# Metabolic engineering of *Corynebacterium glutamicum* for production of the low-caloric natural sweetener D-allulose via phosphorylated intermediates

**DOI:** 10.1101/2025.11.20.689482

**Authors:** Alexander Lehnert, Maja Deditius, Astrid Wirtz, Meike Baumgart, Michael Bott

## Abstract

D-Allulose is a natural, low-calorie sweetener providing ∼70% of the sweetness of sucrose but ≤10% of its caloric value, making it an attractive alternative to conventional sugars. Recently, a phosphorylation–dephosphorylation pathway for D-allulose production was established, involving the formation and irreversible dephosphorylation of D-allulose 6-phosphate. Although this pathway has been demonstrated in *Escherichia coli*, efficient production involved complex medium, complicating downstream processing. Here, we report D-allulose production in minimal medium by implementing the phosphorylation–dephosphorylation pathway in a *Corynebacterium glutamicum* strain unable to metabolize D-fructose. Growth- and production-based screenings identified fructokinase Mak_EC_ and D-allulose 6-phosphate 3-epimerase AlsE_EC_ from *E. coli*, together with D-allulose 6-phosphate phosphatase AlsP_CT_ from *Clostridium thermocellum*, as the most effective enzyme combination for D-allulose formation. Further metabolic engineering of *C. glutamicum* including deletion of *zwf* (D-glucose 6-phosphate dehydrogenase), overexpression of *fbp* (D-fructose 1,6-bisphosphatase), and downregulation of *pgm* (phosphoglucomutase) partially redirected central carbon flux toward D-allulose synthesis, resulting in a 2.3-fold increase in production. The engineered strain produced ∼3.6 g L^−1^ D-allulose from a D-glucose–D-fructose mixture with a yield of 9.1%.

## 1. Introduction

Since the 1970s, obesity has evolved into a global pandemic, which is expected to affect 25% of all adults worldwide by the year 2035 (Lobstein et al., 2024). As driving force for the occurrence of non-communicable diseases (NCDs), which include coronary heart disease, neoplasms, or Diabetes mellitus type 2, obesity is a major public health concern associated with increased mortality and overall decreased life expectancy (Flegal et al., 2013; Fontaine et al., 2003). It is estimated that 10% of the 50.3 million deaths in 2019 were attributable to a too high body mass index (BMI) of ≥25 kg/m^2^ (Lobstein et al., 2024). The rise in obesity is caused by an increased energy intake combined with a reduced energy expenditure, enabled by the greater availability of inexpensive, energy-dense foods and the motorization-related decline in physical activity (Swinburn et al., 2011; Swinburn et al., 2009). A key contributor is caloric sweetener intake, which globally increased by 32% between 1962 and 2000 (Popkin and Nielsen, 2003) and mirrored the rising trend of obesity until the late 1990s (Bray et al., 2004). Although recent trends indicate a decline in sugar intake, overall consumption levels remain high (Malik and Hu, 2022; Newens and Walton, 2016).

Sugarcane- or sugar beet-derived D-sucrose and high-fructose corn syrup (HFCS) still remain the major (high-caloric) sweeteners (OECD/FAO, 2024), which are used for beverages and packaged foods. However, alternative low-caloric sweeteners with little caloric value and a D-sucrose-like sweetness are increasingly gaining interest as substituents, especially for the sweetening of beverages (Sylvetsky and Rother, 2016). The most common alternative sweeteners are saccharin, aspartame, acesulfame-K, sucralose, and sugar alcohols such as sorbitol, mannitol or xylitol. Numerous new sweeteners have been added to the list in recent years, including tagatose and D-allulose. D-Allulose, also known as D-psicose, is a rare sugar with limited availability in nature. It has 70% of the sweetness of D-sucrose but less than 10% (∼0.2 kcal/g) of its caloric content (Moura, 2020). As gel modifier, it can improve food gelling behavior (Ilhan et al., 2020) and since it is a reducing ketohexose, it can undergo the Maillard reaction (Sun et al., 2004), which makes it broadly applicable for baked goods and processed foods. Combined with its attenuation of postprandial blood glucose levels (Franchi et al., 2021) and anti-obesity effects (Chen et al., 2019), D-allulose becomes a suitable candidate for the replacement of high-caloric sweeteners with a wide applicability.

Industrial production of D-allulose involves the utilization of a tagatose 3-epimerase (Itoh et al., 1994) or D-allulose 3-epimerase (Kim et al., 2006), which perform the reversible C3 epimerization of D-fructose to D-allulose at elevated temperatures (55-70 °C) and slightly alkaline conditions (pH 7.5-8.0). The reaction usually stalls at an D-allulose content of 25-35%, making costly separation by simulated moving bed chromatography necessary for increased product yield (Wagner et al., 2015). As an alternative approach, the phosphorylation-dephosphorylation pathway has emerged in recent years, in which the formation and irreversible dephosphorylation of D-allulose 6-phosphate shifts the reaction towards D-allulose formation with a theoretical yield close to 100% (Li et al., 2021). Initially the approach was conducted *in vitro* for D-allulose formation from starch by a multi-enzyme cascade, in which a product yield of 79.2% was obtained (Li et al., 2021). However, purification of in total nine enzymes, addition of phosphorylating agents, and the occurrence of D-glucose and D-fructose as byproducts render the multi-enzyme-based process tedious and expensive for upscaling. Microbial D-allulose production could avoid these problems, as preparation of enzymes and supply of polyphosphate becomes obsolete, while D-glucose and D-fructose formed as byproducts could be reutilized for D-allulose formation. First attempts for microbial D-allulose production were performed with *Escherichia coli* strains using D-fructose (Guo et al., 2022), D-glucose (Taylor et al., 2023), and also D-sucrose as substrates (Zheng et al., 2024). However, these approaches either used complex medium or required the addition of yeast extract for biomass formation, which increases the costs of the overall process and complicates downstream processing. Only in one study, M9 minimal medium was used for the cultivation of an engineered *E. coli* strain to convert D-fructose to D-allulose. However, the strain, which used glycerol as additional carbon source, only formed limited D-allulose titers (1.59 g/L) after 100 h of cultivation (Guo et al., 2022).

*Corynebacterium glutamicum* is a Gram-positive actinobacterium, which has become the major host for the industrial production of amino acids, particularly of L-glutamate and L-lysine (Eggeling and Bott, 2005). In the past decades, its product portfolio was continuously extended and now comprises, e.g. organic acids (Wieschalka et al., 2013), proteins (Freudl, 2017), and high-value active ingredients for food, feed, and human health (Wolf et al., 2021), including rare sugars such as D-tagatose and D-allulose (Jeong et al., 2022; Park et al., 2016). D-Allulose production was performed in biotransformations with intact or permeabilized cells containing D-allulose 3-epimerases that convert D-fructose to D-allulose.

Here, we sucessfully established the phosphorylation-dephosphorylation pathway for D-allulose formation in *C. glutamicum*. For this purpose, we used a strain unable to grow on D-fructose and selected suitable enzymes for synthesis and dephosphorylation of D-allulose 6-phosphate. Subsequently, we engineered central metabolism to increase the carbon flux towards D-allulose.

## 2. Material and methods

### 2.1. Bacterial strains, plasmids and growth conditions

All bacterial strains and plasmids used in this work are listed in Table 1. *E*. *coli* DH5α was used as host for cloning and construction of plasmids. Cultivation of *E. coli* was performed in lysogeny broth (LB) medium or on LB agar plates at 37°C. If the strains carried a plasmid, 50 µg/mL of kanamycin was added to the LB medium or agar. For *C. glutamicum*, cultivations were performed either in brain heart infusion medium (BHI) or in defined CGXII medium (Keilhauer et al., 1993) with 0.03 g/L protocatechuic acid as iron chelator. If the strains carried a plasmid, 25 µg/mL of kanamycin was supplemented to the BHI or CGXII medium.

**Table 1.**
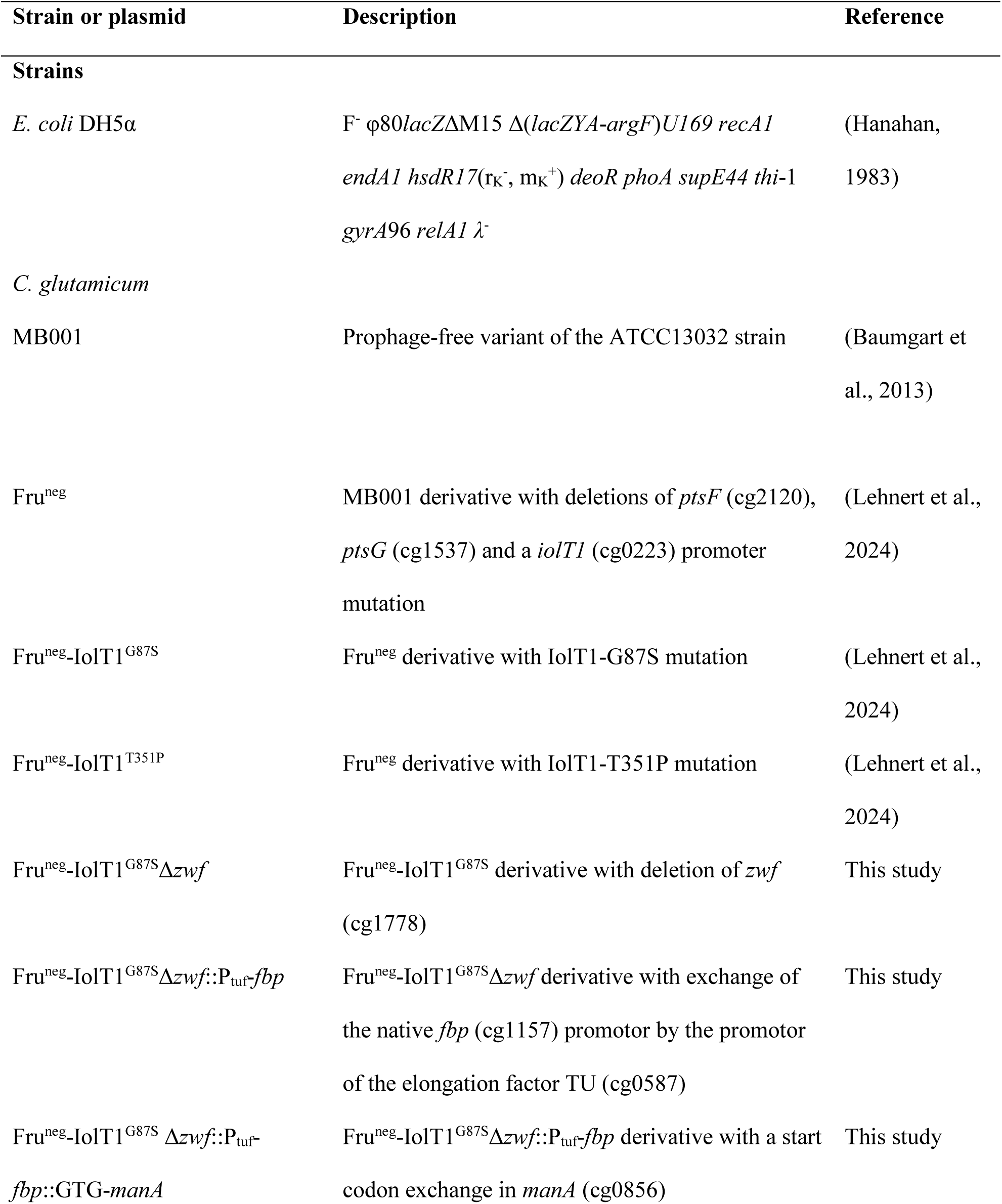
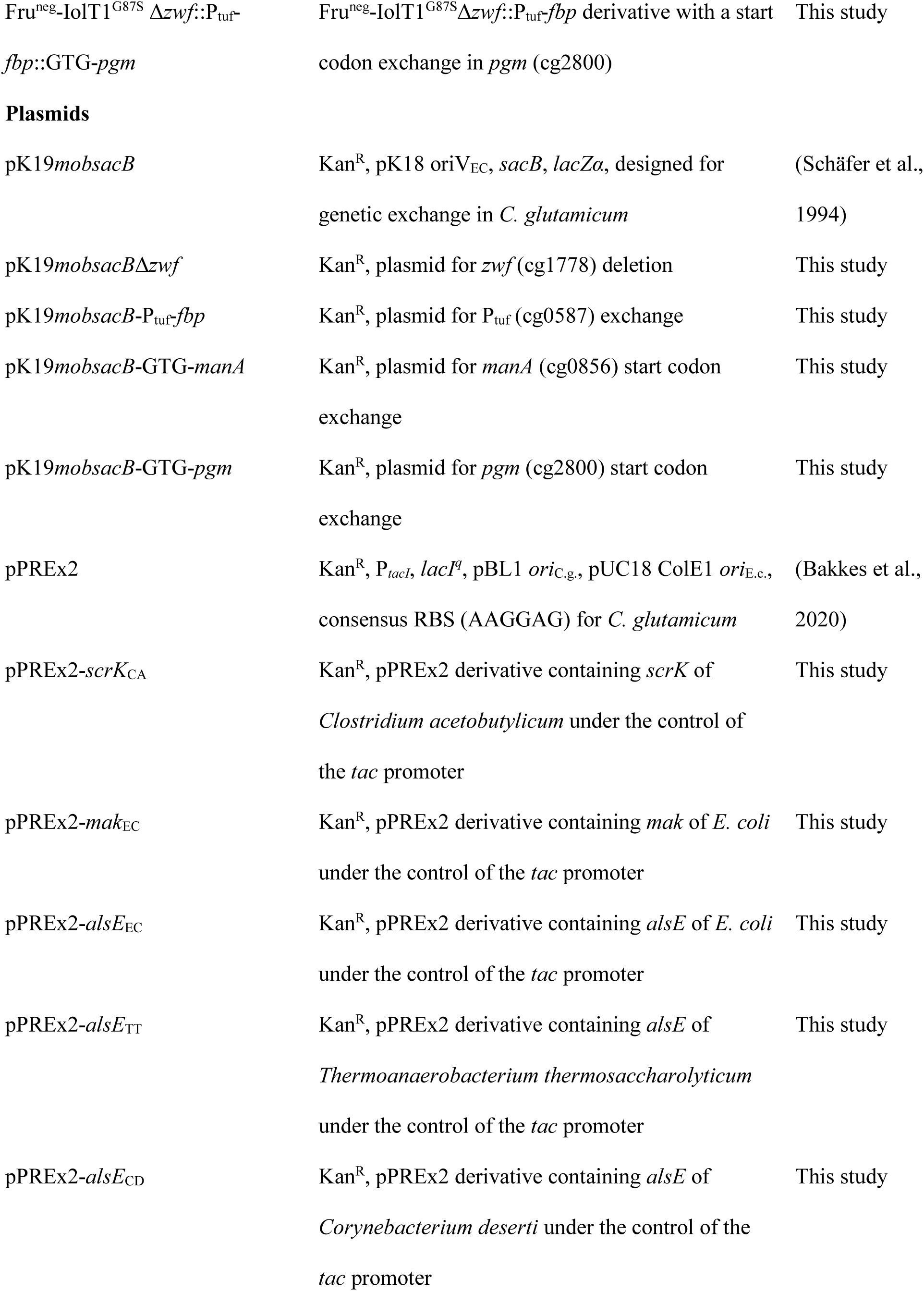
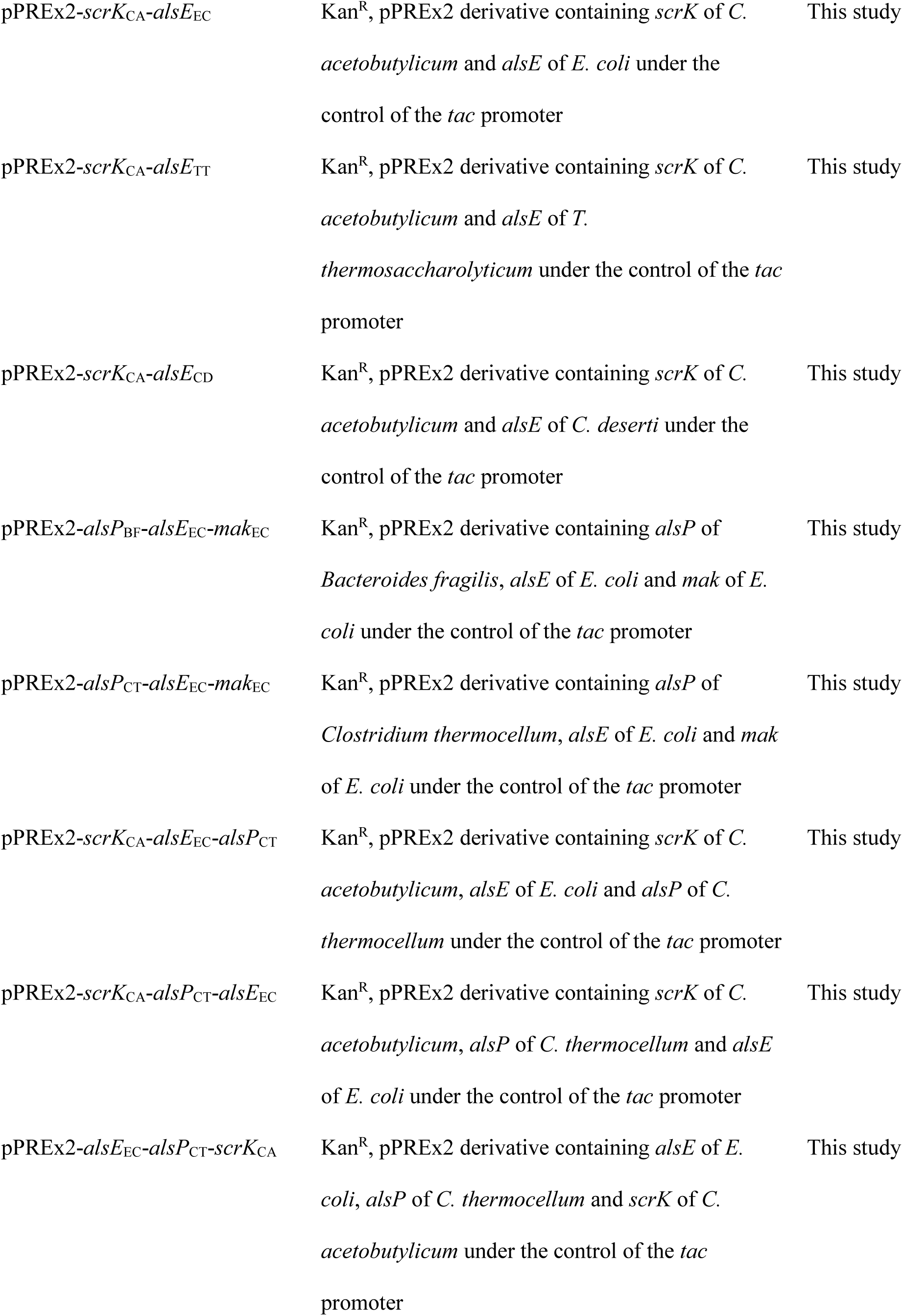
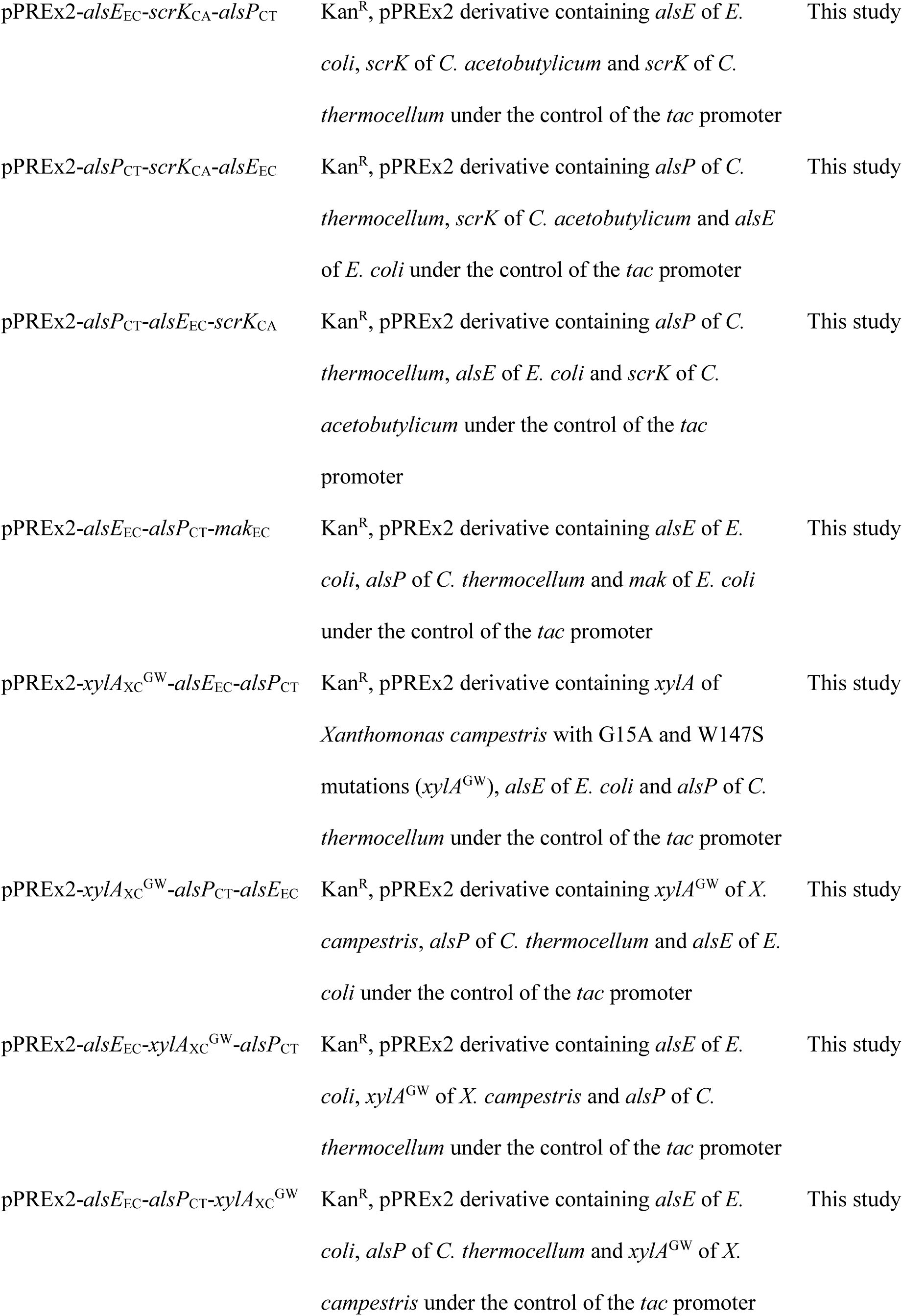
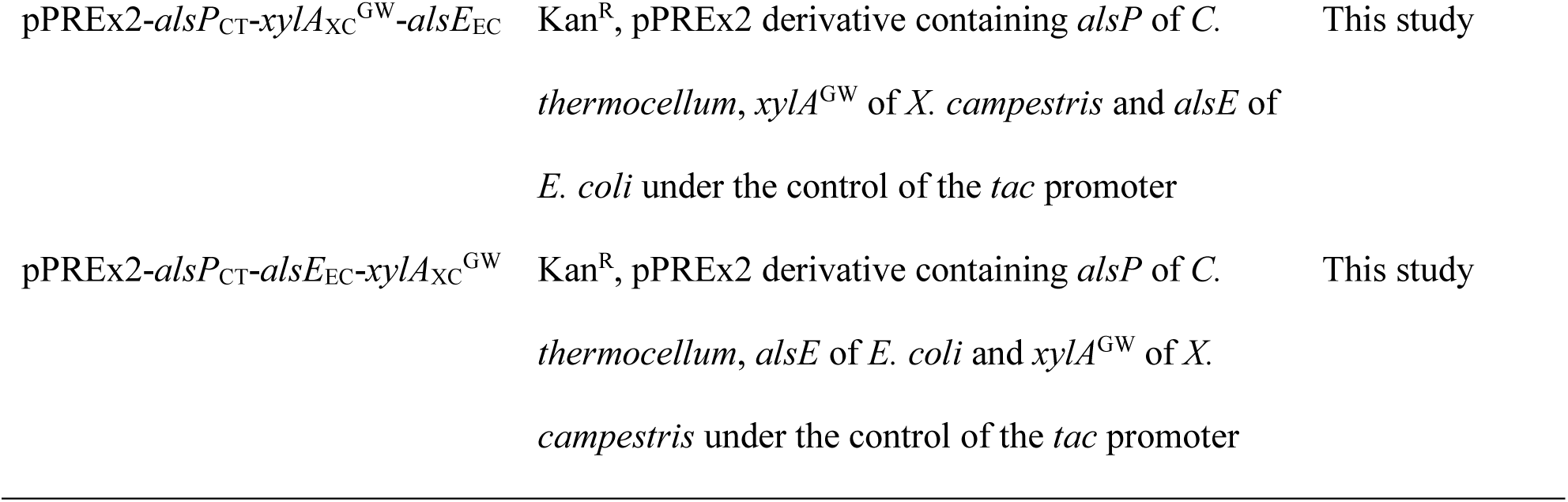
List of strains and plasmids used and constructed during this study.

### 2.2. Recombinant DNA work and construction of deletion mutants

All plasmids and oligonucleotides used in this study are listed in Table 1 and 2, respectively. The fructokinase genes of *Clostridium acetobutylicum* (*scrK*_CA_) and *E. coli* (*mak*_EC_), the D-allulose 6-phosphate 3-epimerase genes of *E. coli* (*alsE*_EC_), *Thermoanaerobacterium thermosaccharolyticum* (*alsE*_TT_) and *Corynebacterium deserti* (*alsE*_CD_), and the D-allulose 6-phosphate phosphatase genes of *Bacteroides fragilis* (*alsP*_BF_) and *Clostridium thermocellum* (*alsP*_CT_) were ordered codon-optimized for *C. glutamicum* from Life Technologies (Carlsbad, USA). PCR, DNA restriction and Gibson assembly were performed according to standard protocols (Gibson et al., 2009; Green and Sambrook, 2012). Transformation of *E. coli* was performed via heat-shock (Hanahan, 1983) and of *C. glutamicum* via electroporation (van der Rest et al., 1999). For *C. glutamicum*, the suicide plasmid pK19*mobsacB* was used for the deletion of genes or the exchange of promoters via double homologous recombination. The pPREx2 plasmid (Bakkes et al., 2020) was used for target gene expression under the control of the isopropyl-β-D-thiogalactoside (IPTG)-inducible *tac* promoter.

**Table 2.**
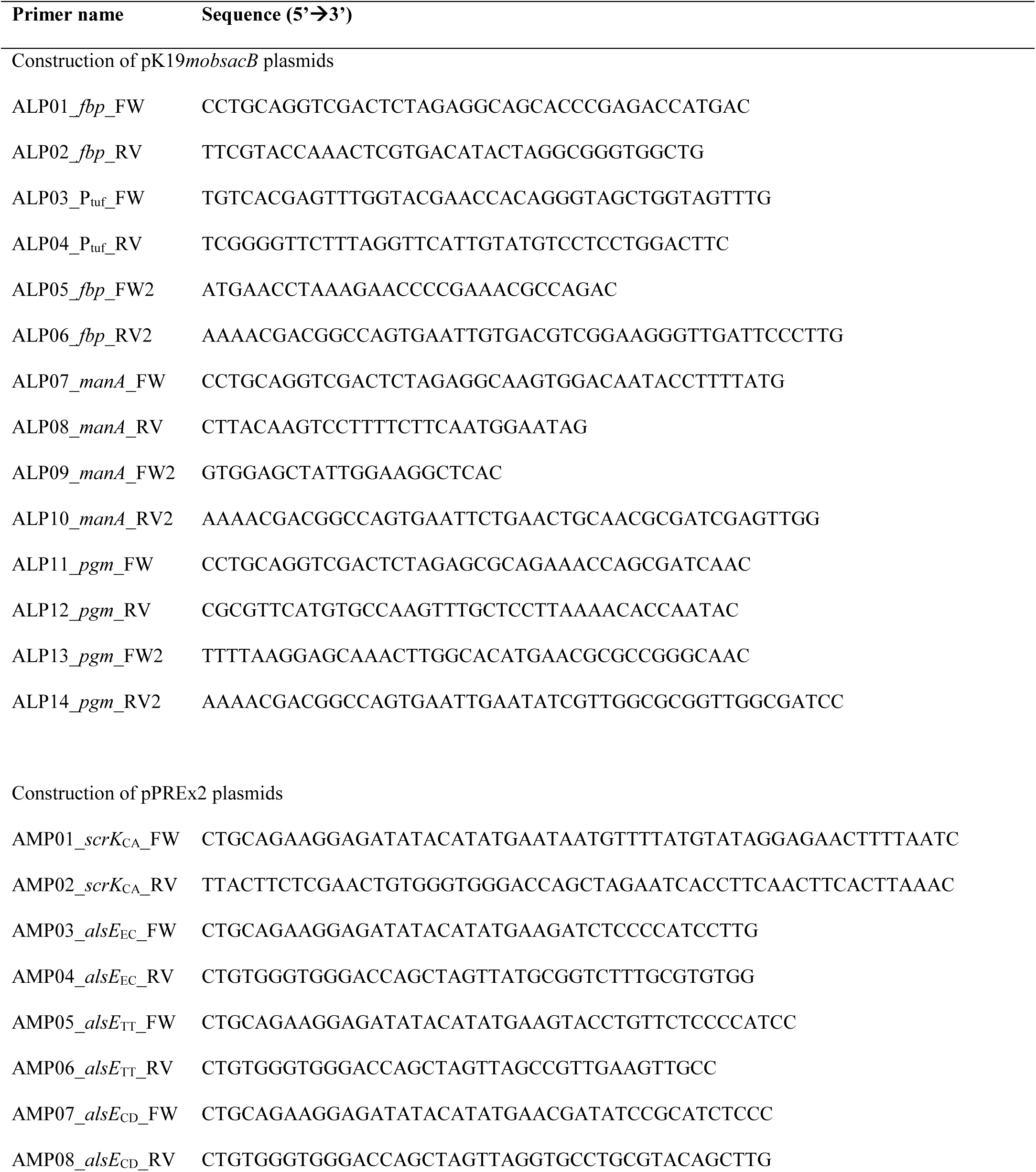
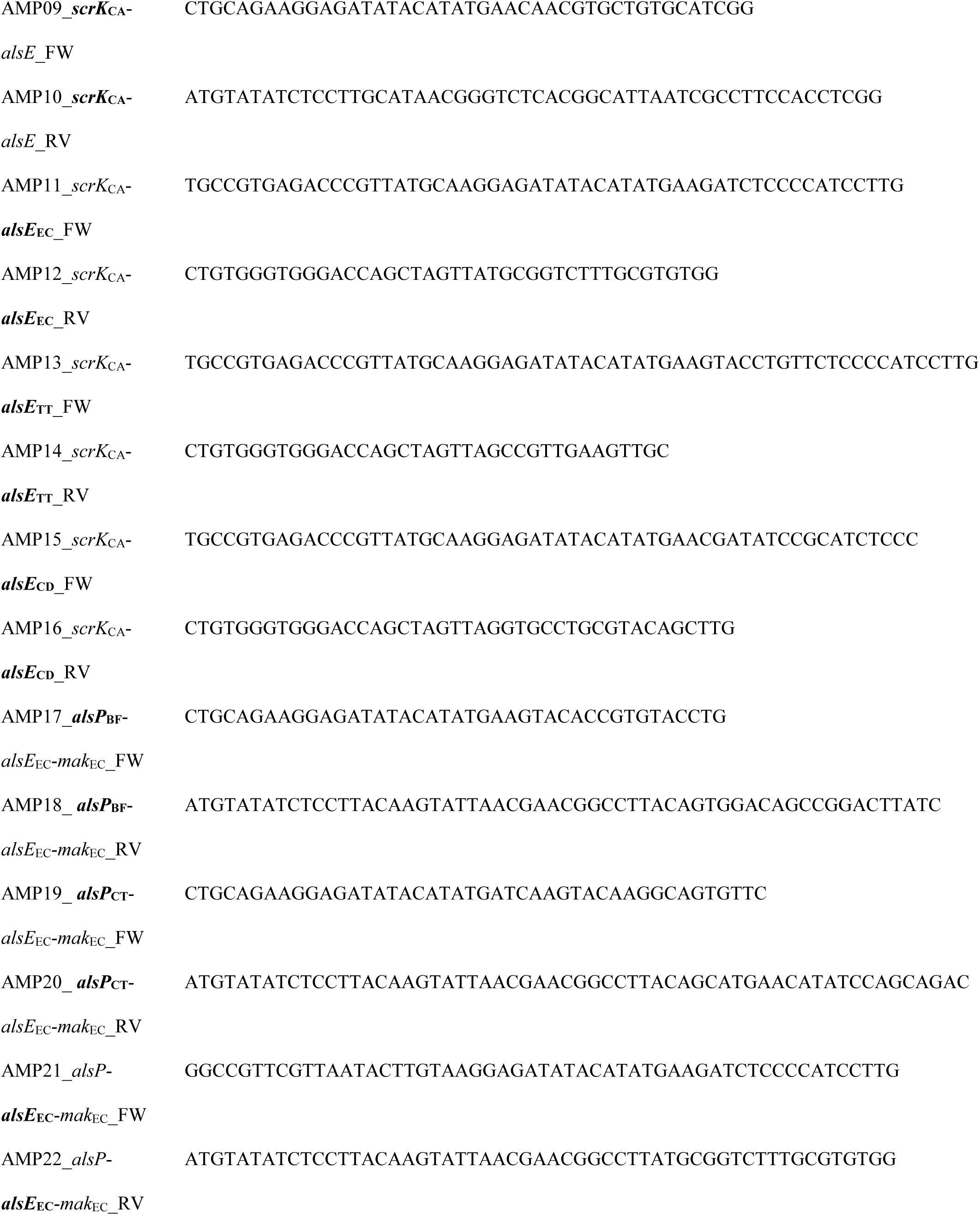
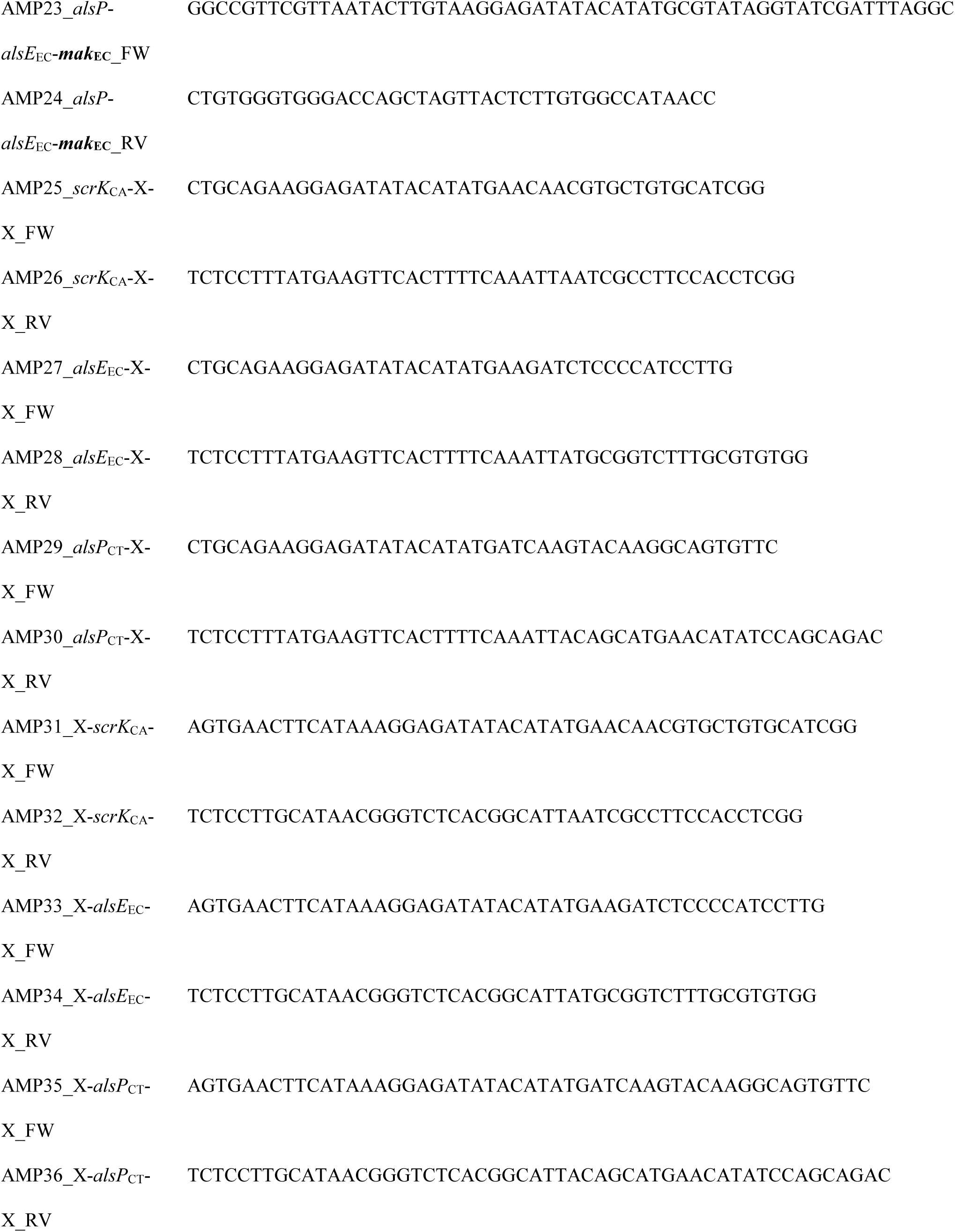
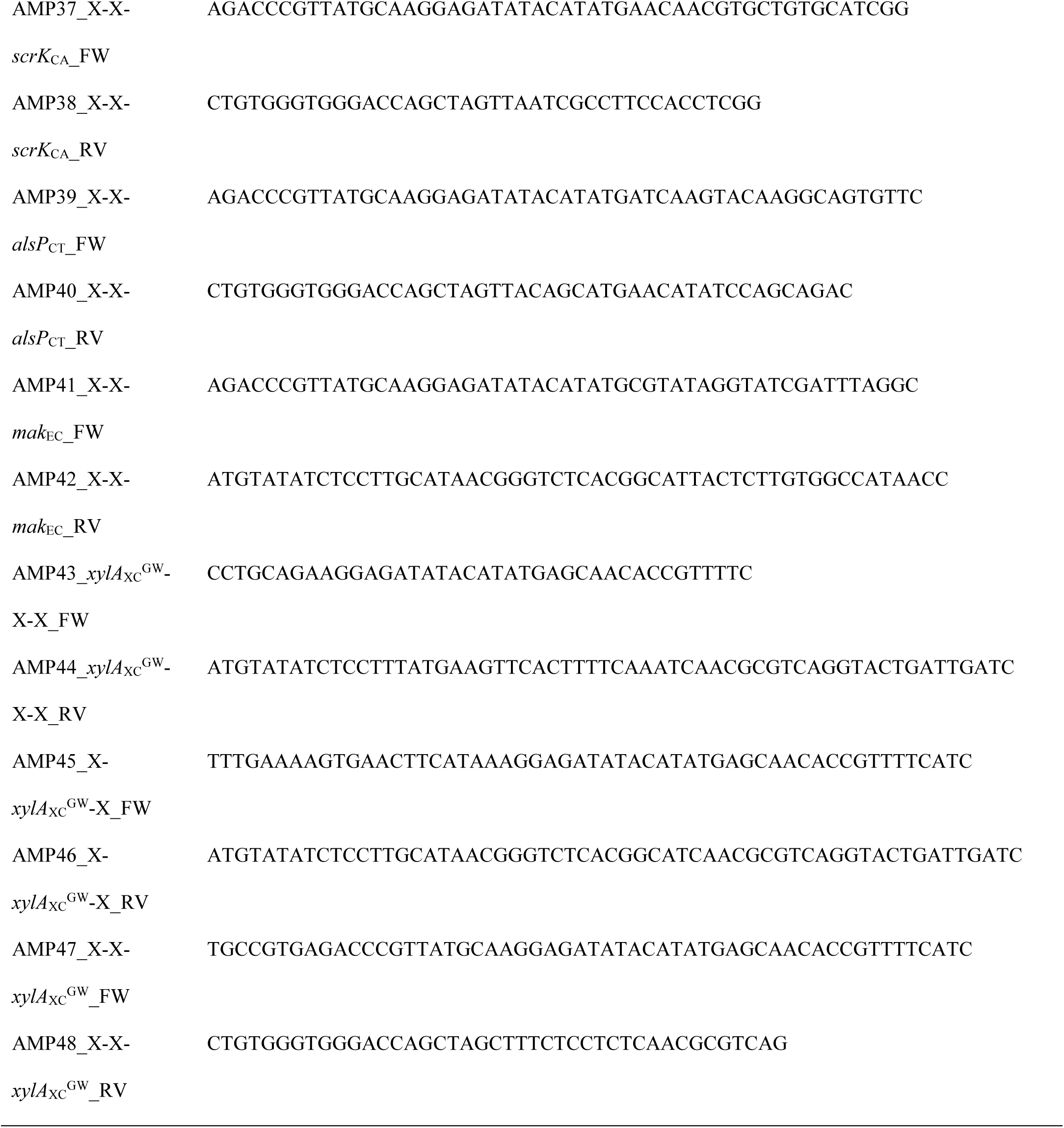
List of oligonucleotides used in this study.

### 2.3. Growth experiments in the BioLector

The BioLector microcultivation system was used for cultivation of *C. glutamicum* strains. It measures growth in 48-well Flowerplates (Beckman Coulter, Brea, USA) by detecting the intensity of backscattered light at 620 nm (Kensy et al., 2009a; Kensy et al., 2009b). All experiments were performed in either the BioLector I or II (Beckman Coulter, Brea, USA) at 30°C, 85% humidity and 1200 rpm. Backscattered light was detected with a signal gain factor of 2 and 4 for the BioLector I and II, respectively. For each cultivation, a FlowerPlate was set up with 800 µL volume of CGXII medium containing either 40 g/L D-fructose or 40 g/L D-allulose with 25 µg/mL kanamycin and 1 mM IPTG. Inoculation was performed to an initial OD_600_ of 5 determined with a spectrophotometer.

### 2.4. D-Allulose production experiments

Production experiments were performed in 500 mL baffled shake flasks at 30°C and 130 rpm with 85% humidity for 72 - 84 h. Two consecutive overnight precultures were prepared before the inoculation of the main cultures. The first precultures were prepared in test tubes with BHI medium containing 25 µg/mL kanamycin that were incubated overnight at 30°C and 170 rpm and used for the inoculation of the second precultures comprising CGXII minimal medium with 20 g/L D-glucose and 25 µg/mL kanamycin. The second precultures were also incubated overnight at 30°C and 170 rpm and then used for the inoculation of the main cultures to an initial OD_600_ of 0.5. Main cultures were prepared in CGXII medium with 20 g/L of D-glucose and 20 g/L of D-fructose, supplemented with 25 µg/mL kanamycin and 1 mM IPTG. Samples of the supernatants were taken after 12 - 24 h, analyzed for growth via OD_600_ measurement and filtered by using 0.2 µm Whatman Puradisc 13 filters (Cytiva, Marlborough, USA). The cell-free filtrate was stored at −20 °C for use in sugar analysis.

### 2.5. Sugar quantification via HPLC

HPLC analysis was conducted with culture supernatant samples diluted 1:4 or 1:8 in deionized water. A Carbo-Pb Guard Cartridge (Phenomenex, Aschaffenburg, Germany) and a Metab-Pb 250 × 7.8 mm column (ISERA, Düren, Germany) were used for the separation of D-glucose, D-fructose, D-sucrose and D-allulose in an Agilent LC-1100 system (Agilent Technologies, Santa Clara, USA). 5 µL of sample was injected into the system and separated at 80°C for 45 min with a flow rate of 0.6 mL/min in double distilled and filtered (0.2 µm filter) water. Detection of sugars was performed with a refraction index detector at 35°C. Calibration curves, obtained from sugar standards with concentrations of 1 g/L, 2.5 g/L, 5 g/L and 10 g/L of D-glucose, D-fructose and D-allulose were used for sugar quantification.

## 3. Results and discussion

### 3.1. Growth-based evaluation of fructokinases

The aim of this study was to establish the production of D-allulose via the phosphorylation-dephosphorylation pathway in *C. glutamicum* from a D-glucose-D-fructose mixture. In *C. glutamicum*, utilization of D-glucose and D-fructose starts with their uptake by the EII permeases of the PEP-dependent phosphotransferase system (PTS). D-Glucose is taken up via PtsG as D-glucose 6-phosphate (Moon et al., 2007), while D-fructose is mainly taken up by PtsF as D-fructose 1-phosphate or, to a much lower extent, by PtsG as D-fructose 6-phosphate (Dominguez and Lindley, 1996; Dominguez et al., 1998). PTS-independent uptake of D-glucose or D-fructose in an unphosphorylated state is also possible by the inositol transporter IolT1 due to its substrate promiscuity (Bäumchen et al., 2009; Brüsseler et al., 2018; Ramp et al., 2022). Once taken up, D-glucose can be phosphorylated by the endogenous glucokinases Glk and PpgK to D-glucose 6-phosphate and further metabolized (Lindner et al., 2010; Park et al., 2000). Since *C. glutamicum* does not possess fructokinase activity, intracellular D-fructose can only be metabolized after export and reuptake by PtsF (Dominguez and Lindley, 1996). Hence, deletion of *ptsF* and *ptsG* is sufficient to abolish D-fructose metabolization in *C. glutamicum* (Moon et al., 2005).

In a recent study, we constructed the fructose-negative *C. glutamicum* strain Fru^neg^, which lacks PtsF and PtsG and constitutively expresses *iolT1* via mutational inactivation of the IolR-repressor binding site in the *iolT1* promotor (Brüsseler et al., 2018). Due to its increased D-fructose import via *iolT1* expression and its inability to metabolize D-fructose, the Fru^neg^ strain was used to screen selected heterologous fructokinases for their ability to enable growth in D-fructose minimal medium (Fig. 1A). Better expression of the fructokinase gene or a higher activity of the respective enzyme itself should result in an increased growth rate in D-fructose minimal medium. Two fructokinases originating from *Clostridium acetobutylicum* (ScrK_CA_) and *Escherichia coli* (Mak_EC_) have so far been functionally expressed in *C. glutamicum* (Hoffmann et al., 2018; Moon et al., 2005). Thus, the corresponding genes were inserted into the expression plasmid pPREx2 and transferred into the Fru^neg^ strain. The recombinant strains were tested for growth in D-fructose minimal medium (Fig. 1A). Surprisingly, neither of the two fructokinases enabled growth in the Fru^neg^ strain (Fig. 1B). Possible explanations could be that the D-fructose uptake rate of IolT1 was too low and/or the intracellular D-fructose concentration might be insufficient to allow a reasonable fructokinase activity required for growth. To test this possibility, we introduced the fructokinase expression plasmids into the strain Fru^neg^-IolT1^T351P^. We recently demonstrated that the T351P amino acid exchange in IolT1 improves D-fructose uptake by at least an order of magnitude (Lehnert et al., 2024). Strain Fru^neg^-IolT1^T351P^ expressing *scrK*_CA_ or *mak*_EC_ was able to grow in D-fructose minimal medium (Fig. 1C). Expression of *scrK*_CA_ resulted in a higher growth rate (µ = 0.388 ± 0.016 h^−1^) and higher endpoint OD_600_ values (36.6 ± 3.9) compared to the expression of *mak*_EC_ (µ = 0.188 ± 0.019 h^−1^; endpoint OD_600_ = 20.0 ± 1.0) (Fig. 1D, Fig. 1E), indicating that ScrK_CA_ performs better than Mak_EC_ in *C. glutamicum*. Therefore, ScrK_CA_ was further used for the identification of the most suitable D-allulose 6-phosphate 3-epimerase.

**Figure 1:**
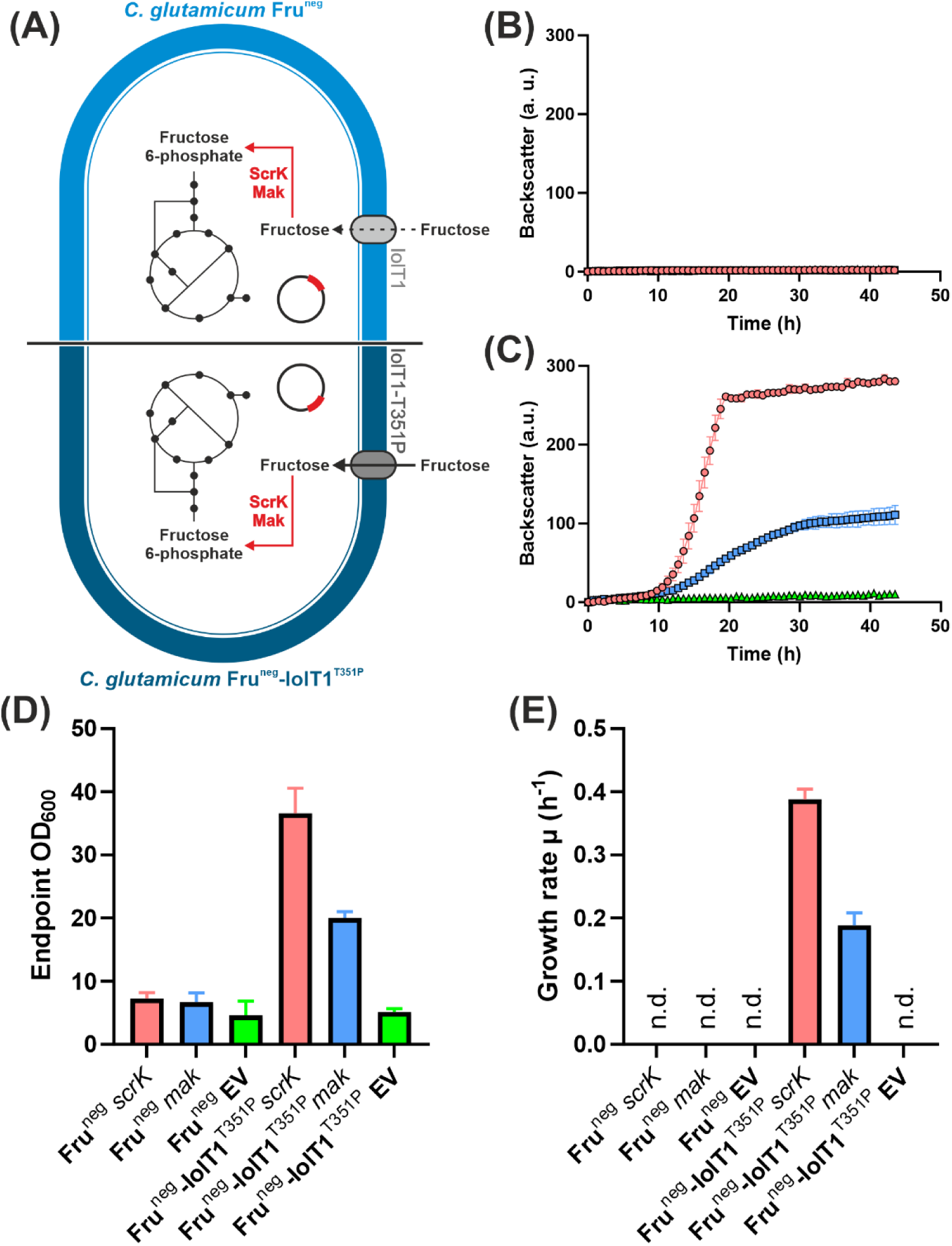
Growth performance in D-fructose minimal medium of the fructose-negative Fru^neg^ strain with either wild-type IolT1 or IolT1-T351P transformed with fructokinase expression plasmids. (A) Scheme of selection strains used for testing the fructokinase genes of *C. acetobutylicum* and *E. coli* provided by the expression plasmids pPREx2-*scrK*_CA_ (red symbols) and pPREx2-*mak*_EC_ (blue symbols). The strains with the empty vector pPREx2 are indicated with green symbols. Growth of Fru^neg^ (B) and Fru^neg^-IolT1^T351P^ (C) carrying either one of the three plasmids in D-fructose minimal medium. (D) Endpoint OD_600_ of the strains after the growth experiment was terminated (43 h). (E) Growth rates obtained for the recombinant Fru^neg^-IolT1^T351P^ strains in a period between 12-16 h of the experiment. The cultivation was performed in CGXII medium with 40 g/L of D-fructose, 1 mM IPTG and 25 µg/mL kanamycin at 30°C and 1200 rpm for 43 h in the BioLector II. Inoculation was performed to an OD_600_ of 5. All data points given represent average values with standard deviations of three biological replicates. N.d. not determinable.

### 3.2. Growth-based evaluation of D-allulose 6-phosphate epimerases

Like for the fructokinase, the most suitable D-allulose 6-phosphate 3-epimerase (AlsE) for *C. glutamicum* was identified via growth experiments. This time, D-fructose was exchanged for D-allulose as sole carbon source in CGXII medium. Simultaneous expression of *scrK*_CA_ and *alsE* should show whether *scrK*_CA_ is able to phosphorylate D-allulose and if so, which of the selected epimerases performs the conversion of D-allulose 6-phosphate to D-fructose 6-phosphate most efficiently, enabling better growth of the strain (Fig. 2A). The D-allulose 6-phosphate 3-epimerases from *E. coli* (AlsE_EC_) and *Thermoanaerobacterium thermosaccharolyticum* (AlsE_TT_) were previously shown to interconvert D-fructose 6-phosphate and D-allulose 6-phosphate (Chan et al., 2008; Li et al., 2021), while the protein from *Corynebacterium deserti* (AlsE_CD_) has only been annotated as a putative D-allulose 6-phosphate 3-epimerase. In each case, the combination of fructokinase and D-allulose 6-phosphate 3-epimerase led to growth of Fru^neg^-IolT1^T351P^ in D-allulose minimal medium (Fig. 2B), suggesting that ScrK_CA_ also accepts D-allulose as substrate. However, lower endpoint OD_600_ values and fourfold decreased growth rates on D-allulose (Fig. 2C, Fig. 2D) compared to D-fructose (Fig. S1) suggest that either uptake of D-allulose and/or phosphorylation by ScrK_CA_ occurred with a much lower efficiency than for D-fructose. The fact that no growth was observed without fructokinase suggests that *C. glutamicum* does not possess an endogenous enzyme activity for the phosphorylation of D-allulose. Of all tested D-allulose 6-phosphate 3-epimerases, AlsE_EC_ enabled the highest growth rate of 0.120 ± 0.007 h^−1^, which is 27% and 37.5% higher compared to the growth rates obtained with AlsE_TT_ (µ = 0.088 ± 0.001 h^−1^) and AlsE_CD_ (µ = 0.075 ± 0.002), respectively (Fig. 2D). The result obtained for AlsE_CD_ confirmed that it can function as D-allulose 6-phosphate 3-epimerase. However, because of the higher growth rate obtained with AlsE_EC_, this enzyme was selected for further growth and D-allulose production experiments.

**Figure 2:**
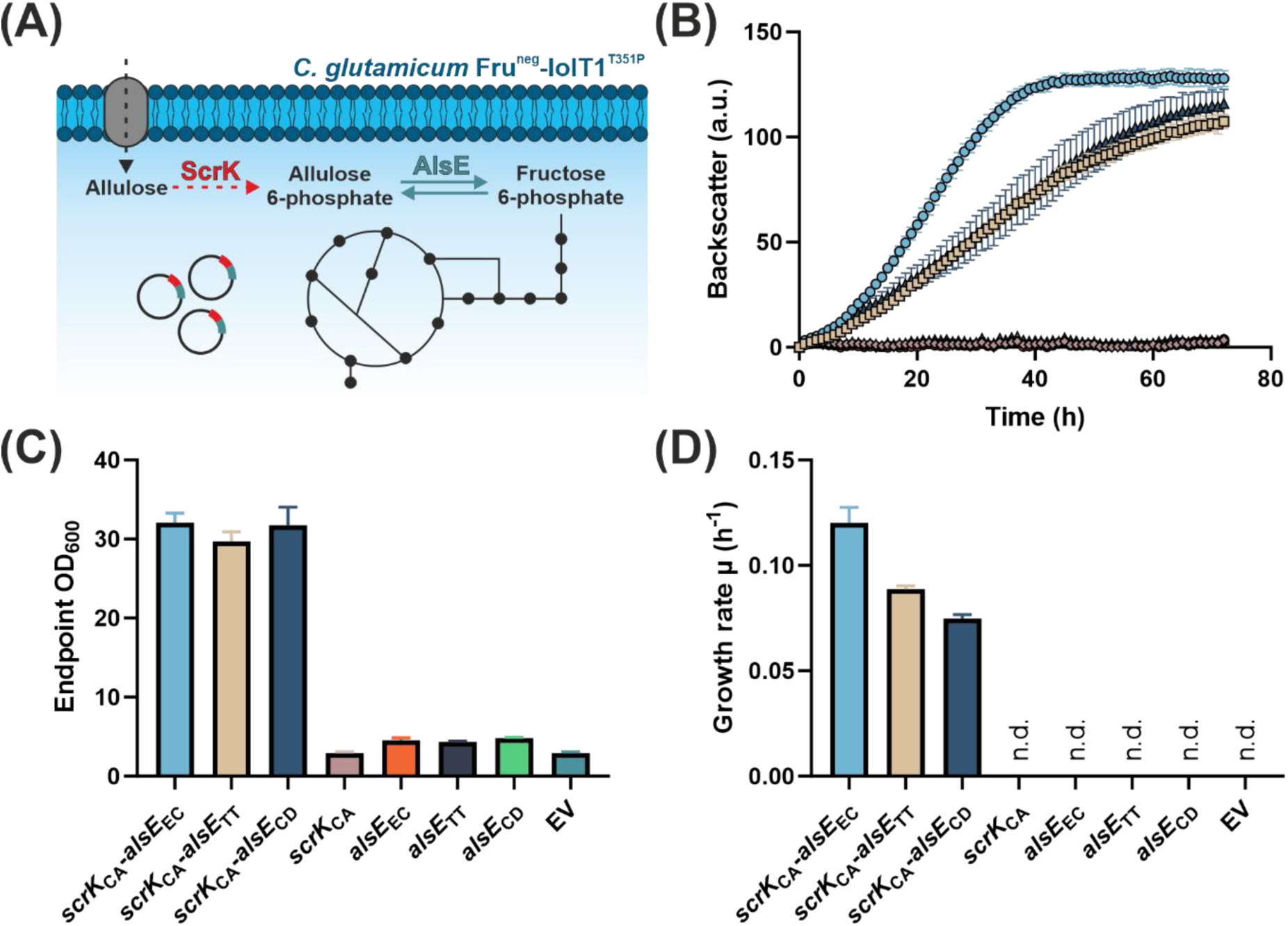
Screening for the most active D-allulose 6-phosphate 3-epimerase in strain Fru^neg^-IolT1^T351P^ via growth in D-allulose minimal medium. (A) Schematic overview of D-allulose metabolism in *C. glutamicum* Fru^neg^-IolT1^T351P^ via plasmid-based expression of the genes for fructokinase (*scrK*) and D-allulose 6-phosphate 3-epimerase (*alsE*). (B) Growth of Fru^neg^-IolT1^T351P^ in CGXII medium with 40 g/L of D-allulose, 1 mM IPTG and 25 µg/mL kanamycin at 30°C and 1200 rpm for 72 h in a BioLector I. The strains were inoculated to an initial OD_600_ of 5. (C) Endpoint OD_600_ obtained after 72 h. (D) Growth rates calculated for each strain in the period between 10-14 h of the experiment. All data points given represent average values with standard deviations of three biological replicates. N.d. not determinable.

### 3.3. Growth- and D-allulose production-based evaluation of D-allulose 6-phosphate phosphatases

In the concept of the phosphorylation-dephosphorylation pathway, irreversibility is achieved by the dephosphorylation of D-allulose 6-phosphate by a phosphatase. The observed phosphorylation of D-allulose by ScrK_CA_ therefore poses a problem, as D-allulose 6-phosphate can be epimerized to D-fructose 6-phosphate and degraded in central metabolism, making the pathway reversible. To prevent this scenario, the activity of the phosphatase should be much higher than that of the kinase phosphorylating D-allulose. Two D-allulose 6-phosphate phosphatases previously described (Li et al., 2021) were tested, the enzymes from *Bacteroides fragilis* (AlsP_BF_) and from *Clostridium thermocellum* (AlsP_CT_). In an initial experiment, the two phosphatases were tested for D-allulose production via plasmid-based expression together with *alsE*_EC_ and *mak*_EC_ in the Fru^neg^ strain (Fig. S2). Although overall D-allulose production was low for both phosphatases, probably due to the limited D-fructose uptake capacity of wild-type IolT1, the strain with AlsP_CT_ reached an almost six times higher final D-allulose titer (0.255 ± 0.037 g/L) compared to the strain with AlsP_BF_ (0.043 ± 0.015 g/L). Therefore, *alsP*_CT_ was selected for the next experiments.

A possibility to vary the ratio between kinase and phosphatase activity is to change the order of their genes with respect to the promoter in the expression plasmid pPREx2. In total six pPREx2-based expression plasmids were constructed carrying the genes *alsP*_CT_, *alsE*_EC_, and *scrK*_CA_ in all possible orders. After transfer into Fru^neg^-IolT1^T351P^, the recombinant strains were cultivated in D-allulose minimal medium and screened for those showing poor growth, which should be due to a high phosphatase/kinase activity ratio (Fig. S3). Three of the six strains showed lower growth rates than the others: pPREx2-*scrK*_CA_-*alsP*_CT_-*alsE*_EC_ (µ = 0.082 ± 0.001 h^−1^), pPREx2-*alsP*_CT_-*alsE*_EC_-*scrK*_CA_ (µ = 0.079 ± 0.001 h^−1^) and pPREx2-*alsE*_EC_-*alsP*_CT_-*scrK*_CA_ (µ = 0.078 ± 0.006 h^−1^). The corresponding strains were then tested for D-allulose production in CGXII medium with 20 g/L of D-glucose and 20 g/L of D-fructose (Fig. S4). During cultivation, D-allulose production peaked at 24 h, reaching titers between 1.2 – 1.9 g/L of D-allulose. Afterwards the D-allulose content declined continuously, suggesting that the dephosphorylation activity provided by AlsP_CT_ is not sufficient to prevent D-allulose rephosphorylation by ScrK_CA_. After 72 h, the strain with pPREx2-*alsE*_EC_-*alsP*_CT_-*scrK*_CA_ showed the highest residual D-allulose titer of 0.490 ± 0.093 g/L, followed by pPREx2-*alsP*_CT_-*alsE*_EC_-*scrK*_CA_ (0.390 ± 0.141 g/L) and pPREx2-*scrK*_CA_-*alsP*_CT_-*alsE*_EC_ (0.360 ± 0.100 g/L) (Fig. S4).

Based on these results, we exchanged *scrK*_CA_ by *mak*_EC_ (Fig. 3A), as Mak_EC_ in strain Fru^neg^-IolT1^T351P^ led to a much lower growth rate on D-fructose than ScrK_CA_ (Fig. 1). Similarly, when provided together with AlsE_EC_, Mak_EC_ led to a lower growth rate on D-allulose (µ = 0.088 ± 0.009 h^−1^) than ScrK_CA_ (µ = 0.167 ± 0.003 h^−1^) (Fig. 3B, C). The additional presence of AlsP_CT_ phosphatase activity further reduced these growth rates (Fig. 3B, C). The Fru^neg^-IolT1^T351P^ strain carrying pPREx2-*alsE*_EC_-*alsP*_CT_-*mak*_EC_ revealed the lowest growth rate with 0.067 ± 0.006 h^−1^ (Fig. 3C) and the replacement of *scrK*_CA_ by *mak*_EC_ enabled an increased and stabilized D-allulose production, which was almost fourfold higher compared to the strain with *alsE*_EC_-*alsP*_CT_-*scrK*_CA_ (Fig. S4) and reached a final titer of 1.904 ± 0.341 g/L (Fig. 3D).

**Figure 3:**
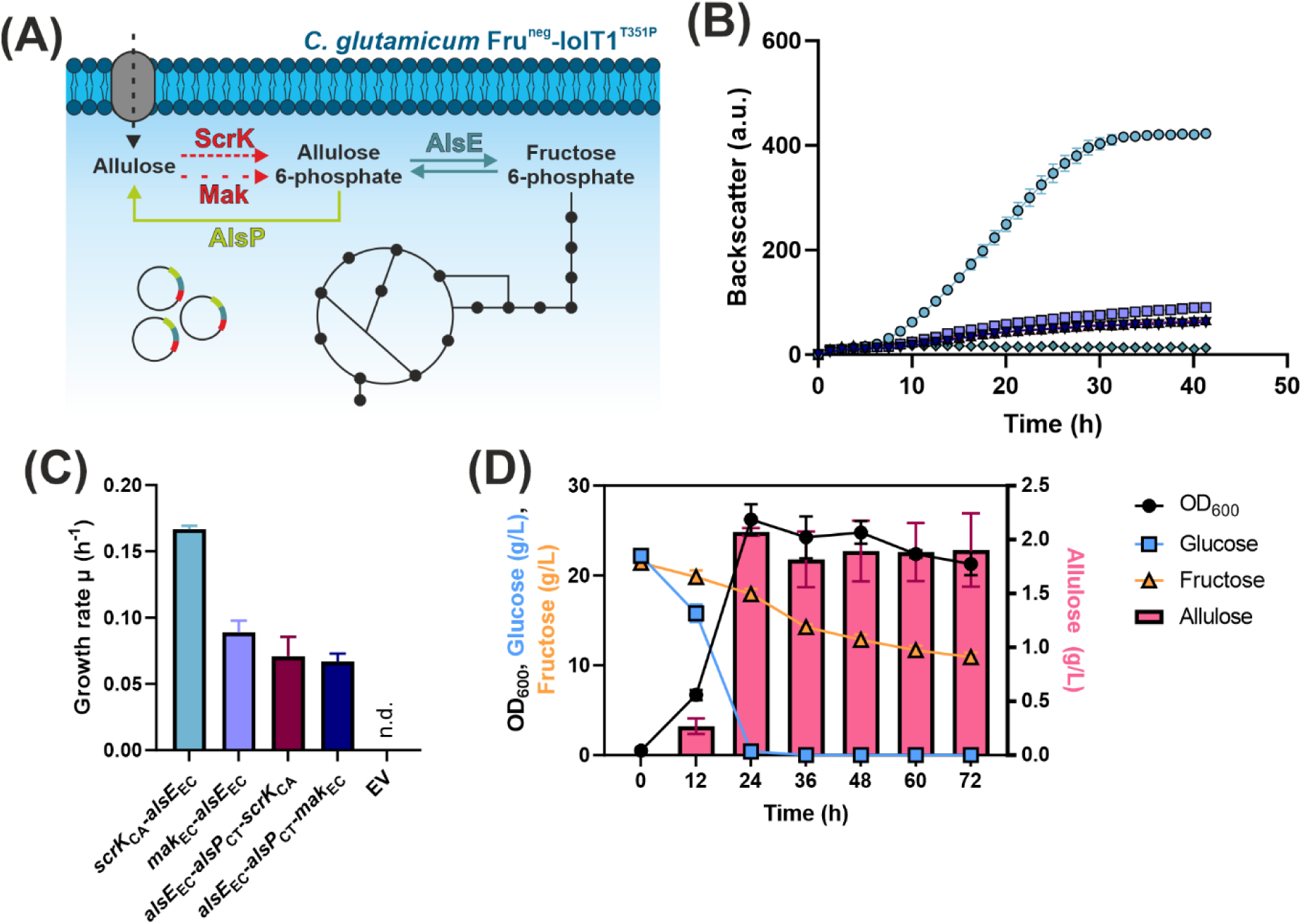
Influence of Mak_EC_ on growth with D-allulose and on D-allulose production in strain Fru^neg^-IolT1^T351P^. (A) Overview of D-allulose metabolization in *C. glutamicum* with ScrK, Mak, AlsE and AlsP. (B) Growth experiment with strains expressing the genes indicated in panel C in CGXII medium with 40 g/L of D-allulose at 30°C and 1200 rpm in a BioLector I for 42 h. Inoculation was performed to an initial OD_600_ of 5. (C) Growth rates of the strains cultivated in D-allulose minimal medium. For the strains carrying pPREx2-*scrK*_CA_-*alsE*_EC_ and pPREx2-*mak*_EC_-*alsE*_EC,_ the period between 8.34 to 11 h was selected for growth rate calculation, while the period between 12.67 to 15 h was used for all other strains. (D) Production experiment of strain Fru^neg^-IolT1^T351P^ pPREx2-*alsE*_EC_-*alsP*_CT_-*mak*_EC_ in CGXII medium with 20 g/L of D-glucose, 20 g/L of D-fructose, 1 mM IPTG and 25 µg/mL kanamycin. The experiment was performed at 30°C and 130 rpm for 72 h. All data points given represent average values with standard deviations of three biological replicates. N.d. not determinable.

### 3.4. The xylose isomerase-based approach

As an alternative approach that avoids the presence of a fructokinase and thus the phosphorylation of D-allulose, we tested the use of a xylose isomerase converting D-fructose to D-glucose, which is then phosphorylated by the endogenous kinases Glk and PpgK to D-glucose 6-phosphate and converted to D-fructose 6-phosphate by D-glucose 6-phosphate isomerase (Pgi). For this purpose, a recently evolved xylose isomerase from *Xanthomonas campestris* was used that carried the amino acid exchanges G15A and W147S (XylA_XC_^GW^) and showed increased D-glucose-D-fructose conversion activity at 30°C (Lehnert et al., 2024). The fructose kinase gene was replaced by the *xylA*_XC_^GW^ gene in the pPREx2-based expression plasmids containing also *alsE*_EC_ and *alsP*_CT_. As before, six plasmids with the genes in all possible orders were constructed and tested in strain Fru^neg^-IolT1^T351P^ for D-allulose production from D-glucose and D-fructose (Fig. 4). All strains reached the highest D-allulose titer (up to 2 g/L) after 24-36 h of cultivation, when D-glucose had been completely consumed. However, afterwards the D-allulose concentration decreased continuously until the cultivation was terminated after 72 h. This means that despite the absence of a fructokinase, D-allulose can be degraded in these strains. The most likely explanation is given by the observation that D-allulose 6-phosphate epimerases can also interconvert the unphosphorylated sugar with low efficiency (Li et al., 2021). As a consequence, D-allulose is converted to D-fructose, which then is isomerized by XylA_XC_^GW^ to D-glucose. D-Glucose is then phosphorylated and converted to D-fructose 6-phosphate, which will enter glycolysis, but can also be converted to D-allulose again. Therefore, further metabolic engineering strategies were tested to improve D-allulose synthesis and avoid its degradation.

**Figure 4:**
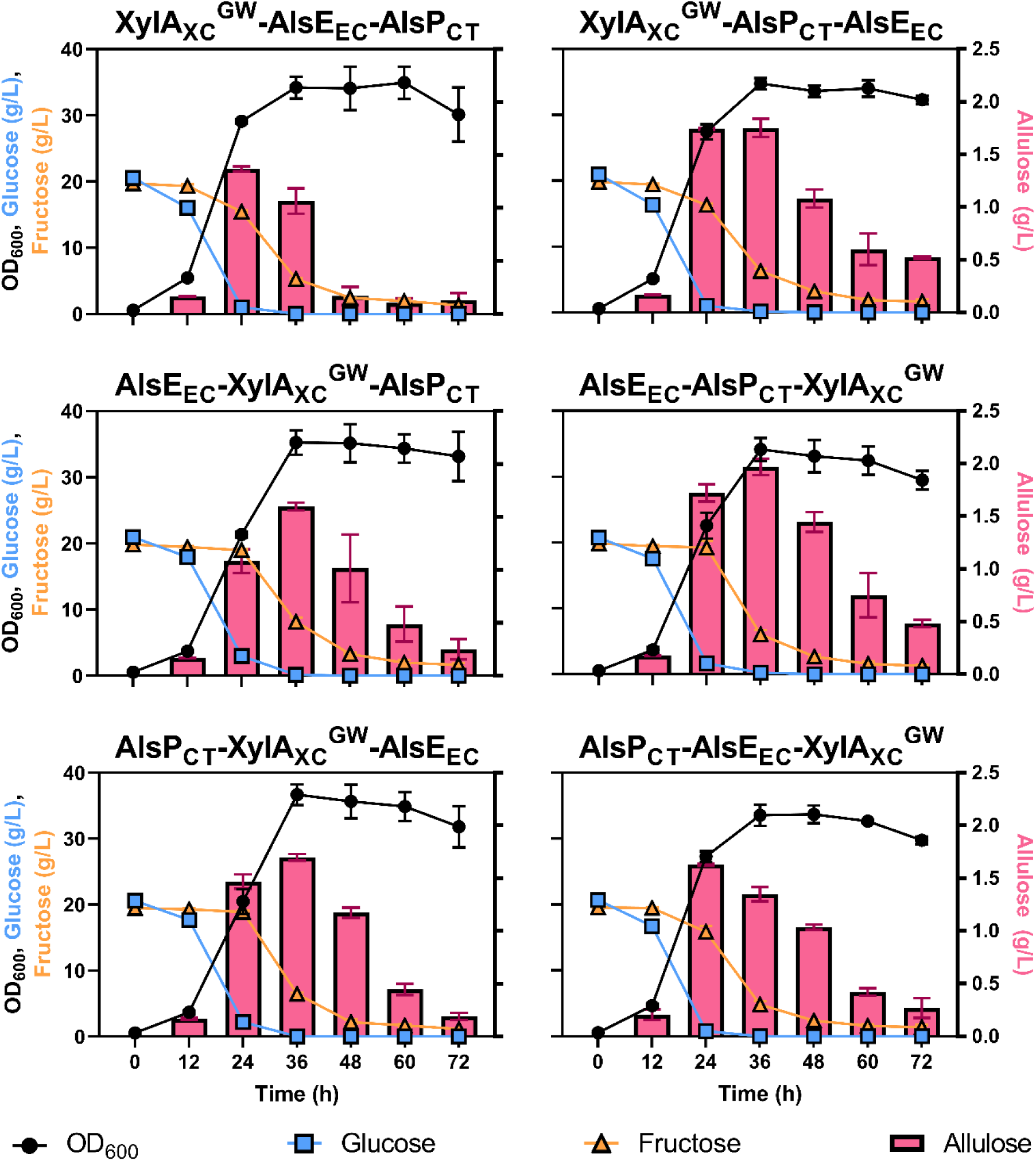
Xylose isomerase mediated production of D-allulose. The Fru^neg^-IolT1^T351P^ strain, transformed with pPREx2 derivatives encoding the three enzymes listed above each panel in the corresponding order, was used for cultivation. The culvation was performed in CGXII medium with 20 g/L of D-glucose, 20 g/L of D-fructose, 1 mM IPTG and 25 µg/mL kanamycin at 30°C and 130 rpm for 72 h. Samples of the supernatant were taken every 12 h. Growth was measured via OD_600_ and sugar concentrations via HPLC. The data points represent average values with standard deviations of three biological replicates.

### 3.5. Slowing down substrate uptake to stabilize product formation

We assume that D-allulose is taken up by *C. glutamicum* via IolT1. The T351P mutation in IolT1 was previously shown to boost uptake of D-glucose and D-fructose into *C. glutamicum* and might also do so for D-allulose. In this case, after consumption of the substrates D-glucose and D-fructose, D-allulose will be taken up into the cell and degraded either by direct phosphorylation or after conversion to D-fructose. In order to potentially reduce D-allulose uptake, IolT1-T351P was replaced by IolT1-G87S, which is characterized by a somewhat lower D-glucose and D-fructose uptake (Lehnert et al., 2024). Therefore, plasmids pPREx2-*alsE*_EC_-*alsP*_CT_-*mak*_EC_ and pPREx2-*alsE*_EC_-*alsP*_CT_-*xylA*_XC_^GW^ were transferred into strain Fru^neg^-IolT1^G87S^ and tested for D-allulose production from D-glucose and D-fructose (Fig. 5). The fructokinase-based approach was barely affected by IolT1^G87S^, as the progression of D-allulose formation mimicked the production in the Fru^neg^-IolT1^T351P^ strain (Fig. 3D, Fig. 5). The highest titer of 2.045 ± 0.062 g/L was reached after 24 h and decreased thereafter to 1.833 ± 0.048 g/L after 72 h (Fig. 5). For the xylose isomerase-based approach, the introduction of IolT1^G87S^ stabilized D-allulose production. Although the pattern was similar to the Fru^neg^-IolT1^T351P^ strain (Fig. 4), which peaked at 36 h and decreased afterwards, D-allulose degradation in the Fru^neg^-IolT1^G87S^ strain was less severe. After 72 h, the culture of the Fru^neg^-IolT1^G87S^ strain still contained 0.822 ± 0.031 g/L of D-allulose, which is 1.7-times higher than the final titer of the Fru^neg^-IolT1^T351P^ strain (0.482 ± 0.035 g/L). Based on these results, the Fru^neg^-IolT1^G87S^ strain was chosen in further efforts for strain improvement.

**Figure 5:**
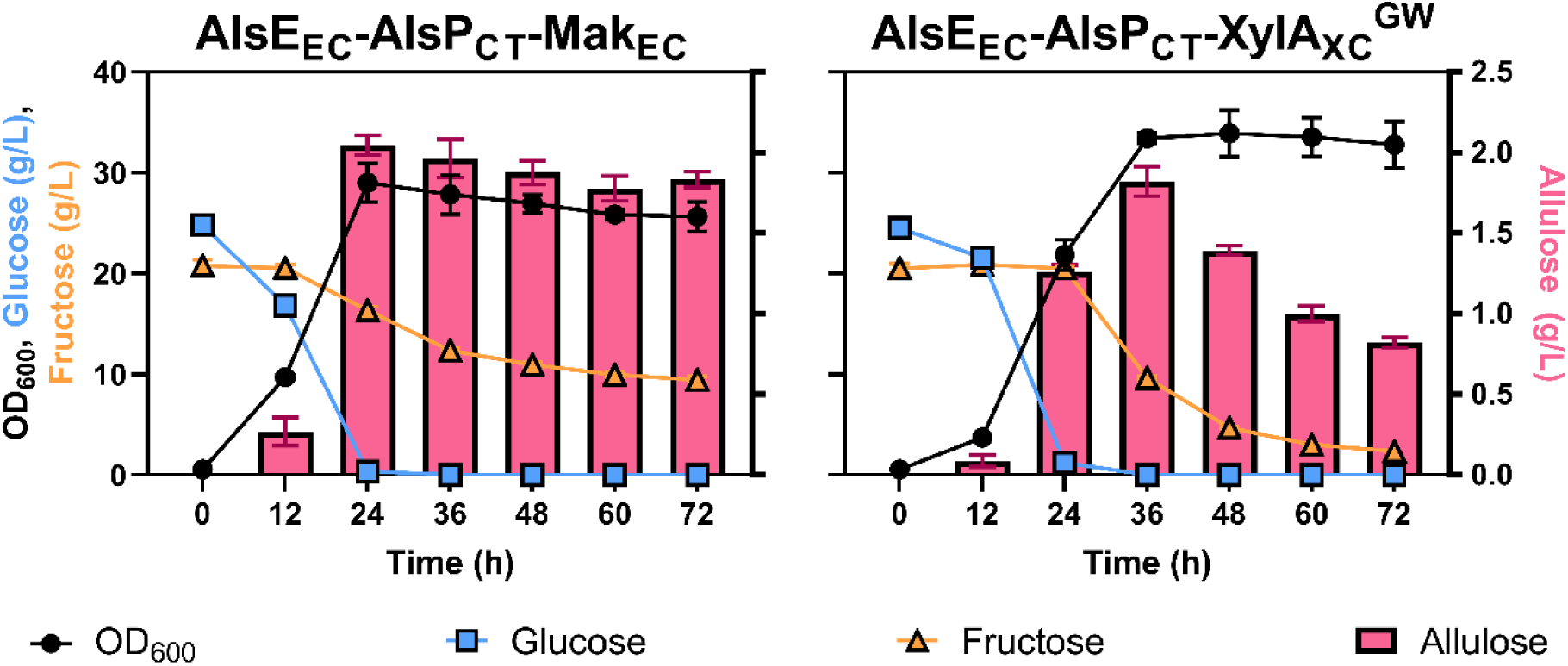
Influence of IolT1-G87S on D-allulose formation. The Fru^neg^-IolT1^G87S^ strain, transformed with pPREx2-*alsE*_EC_-*alsP*_CT_-*mak*_EC_ (AlsE_EC_-AlsP_CT_-Mak_EC_) or pPREx2-*alsE*_EC_-*alsP*_CT_-*xylA*_XC_^GW^ (AlsE_EC_-AlsP_CT_-XylA_XC_^GW^) was cultivated in CGXII medium with 20 g/L of D-glucose and 20 g/L of D-fructose at 30°C and 130 rpm for 72 h. In 12 h intervals, samples were taken. Growth was determined via OD_600_ measurement and sugar concentrations in the supernatant were determined via HPLC. The data points represent average values with standard deviations of three biological replicates.

### 3.6. Metabolic engineering of central metabolism for increased D-fructose 6-phosphate supply

Independent of whether D-allulose is formed via the fructokinase- or the xylose isomerase-based phosphorylation-dephosphorylation pathway, it is hampered by the fact that D-glucose 6-phosphate and D-fructose 6-phosphate are intermediates of central carbon metabolism and thus offer a variety of possibilities to escape D-allulose formation (Fig. 6). To increase the carbon flux towards D-allulose, the *zwf* gene encoding D-glucose 6-phosphate dehydrogenase (Moritz et al., 2000) was deleted in strain Fru^neg^-IolT1^G87S^, which was then transformed with pPREx2*-alsE*_EC_*-alsP*_CT_*-mak*_EC_ and pPREx2*-alsE*_EC_*-alsP*_CT_*-xylA*_XC_^GW^. The absence of *zwf* caused reduced growth and lowered D-glucose and D-fructose consumption rates (Fig. S5). D-Allulose yields slightly decreased for the fructokinase-expressing strain (3.8 ± 0.4%), while they almost tripled for the xylose isomerase-expressing strain (5.2 ± 0.2%) after 72 h (Fig. S5, Fig. 7A). However, since growth was negatively affected by the *zwf* deletion causing a reduction of the endpoint OD_600_ by 22% for the fructokinase-expressing strain (OD_600_ = 19.9 ± 1.4) and of 30% for the xylose isomerase-expressing strain (OD_600_ = 23.1 ± 2.2), the OD_600_-normalized D-allulose production increased for both Δ*zwf* strains compared to the *zwf*-containing parent strains (Fig. S5). The positive effect of the *zwf* deletion on xylose isomerase-based D-allulose production can be explained by a decelerated degradation of D-glucose 6-phosphate. Since D-glucose 6-phosphate is the precursor for xylose isomerase-based D-allulose formation, avoiding its oxidation to 6-phosphogluconate allows a higher conversion to D-allulose.

**Figure 6:**
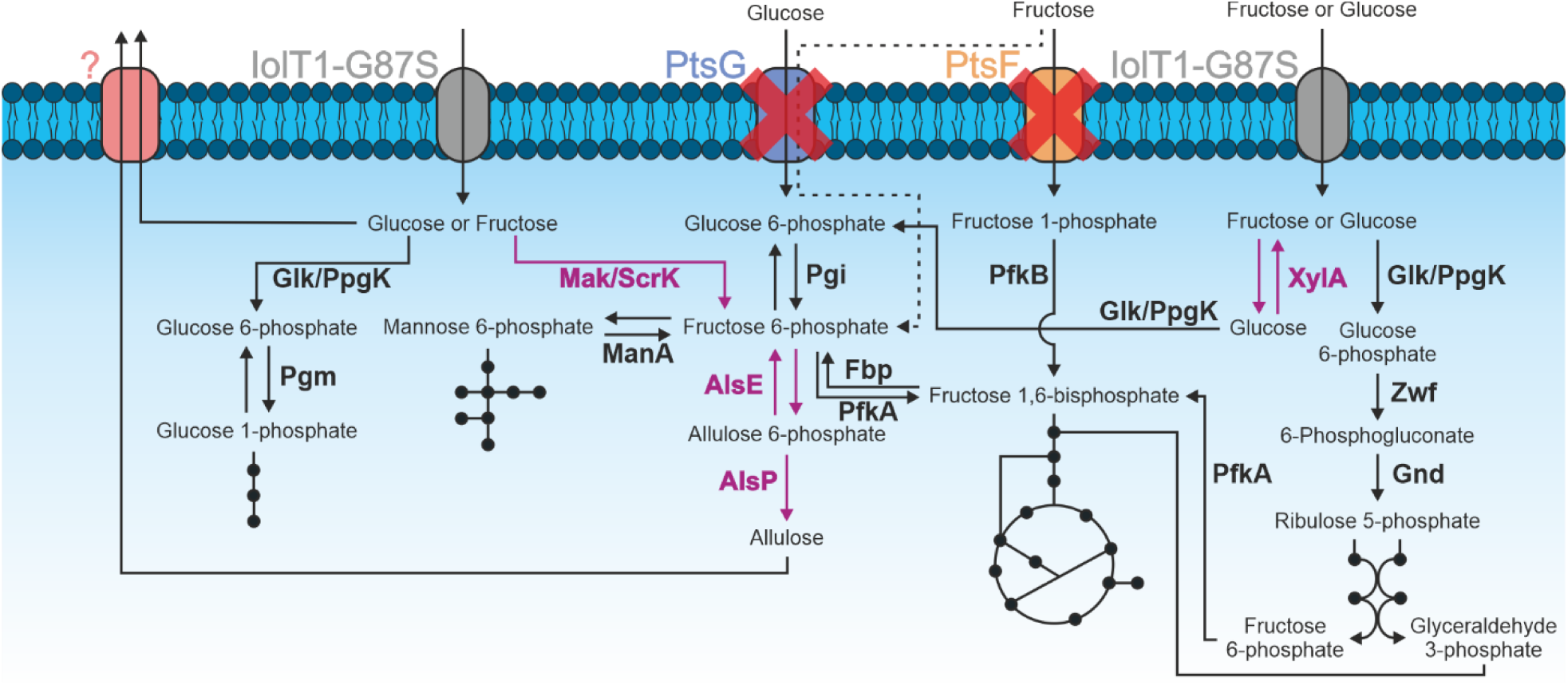
Sugar metabolism and D-allulose production in *C. glutamicum* Fru^neg^-IolT1^G87S^. The black arrows illustrate endogenous reactions in *C. glutamicum*. The violet arrows indicate reactions catalyzed by heterologous enzymes. Proteins, which have been deleted are marked with a red cross. Abbreviations: AlsE, D-allulose 6-phosphate 3-epimerase; AlsP, D-allulose 6-phosphate phosphatase; Fbp, D-fructose 1,6-bisphosphatase; Glk, ATP-dependent glucokinase; Gnd, 6-phosphogluconate dehydrogenase; IolT1-G87S, inositol transporter with G87S mutation for improved D-glucose and D-fructose uptake; Mak, fructokinase of *E. coli*; ManA, D-mannose 6-phosphate isomerase; PfkA, 6-phosphofructokinase; PfkB, 1-phosphofructokinase; Pgi, phosphoglucoisomerase; Pgm, phosphoglucomutase; PpgK, polyphosphate-dependent glucokinase; PtsF, EII permease for the uptake of D-fructose; PtsG, EII permease for the uptake of D-glucose; ScrK, fructokinase of *C. acetobutylicum*; XylA, xylose isomerase from *X. campestris* with two amino acid exchanges; Zwf, D-glucose 6-phosphate dehydrogenase; ?, unknown D-fructose/D-allulose exporter.

**Figure 7:**
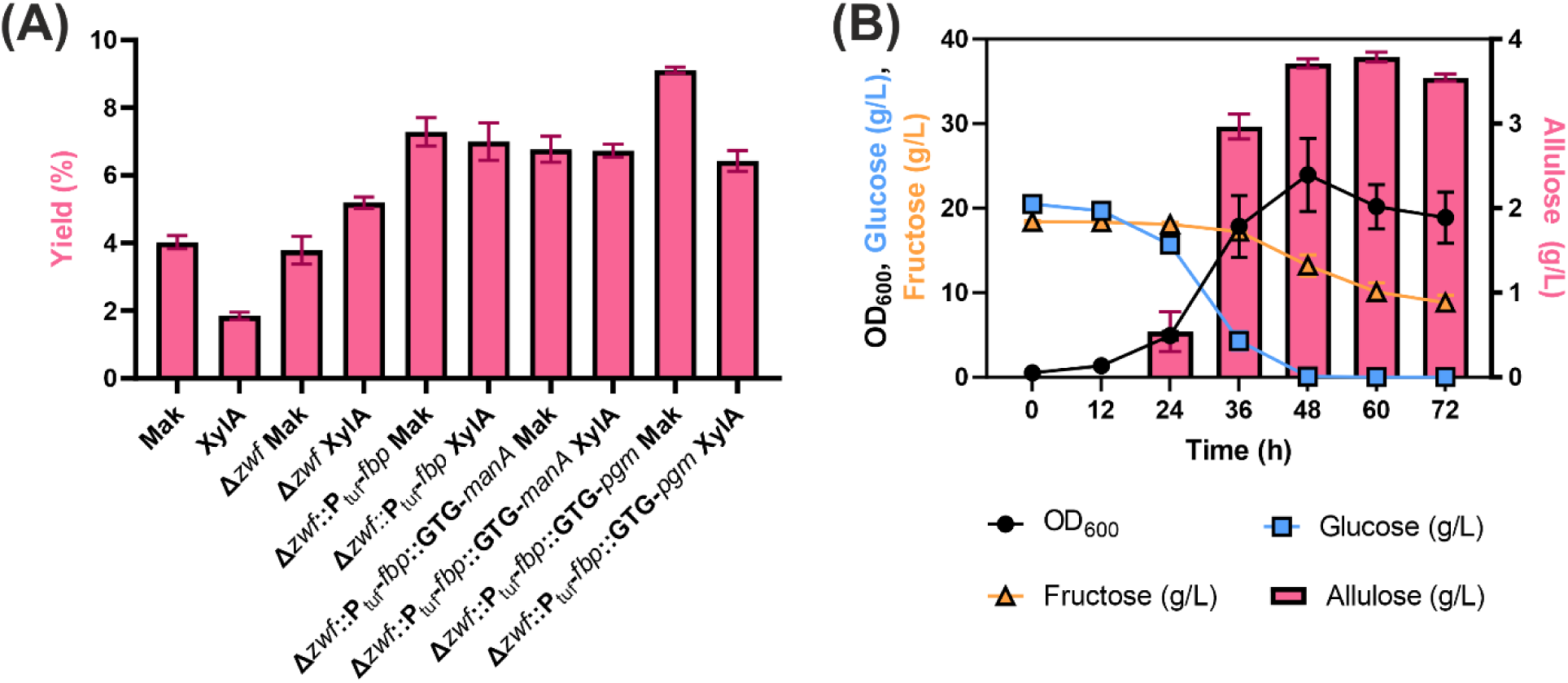
Influence of metabolic engineering approaches in *C. glutamicum* Fru^neg^-IolT1^G87S^ on the yield of D-allulose from D-glucose and D-fructose. The strains were transformed either with pPREx2-*alsE*_EC_-*alsP*_CT_-*mak*_EC_ (Mak) or with pPREx2-*alsE*_EC_-*alsP*_CT_-*xylA*_XC_^GW^ (XylA). A D-glucose 6-phosphate dehydrogenase deletion mutant (Δ*zwf*), a Δ*zwf* mutant with a *tuf* promotor exchange of the D-fructose 1,6-bisphosphatase (P_tuf_-*fbp*), and a Δ*zwf*::P_tuf_-*fbp* mutant with either a start codon exchange of the mannose 6-phosphate isomerase (GTG-*manA*) or the phosphoglucomutase (GTG-*pgm*) was used. (A) D-Allulose yields of the respective strains transformed with the either the Mak or the XylA expressing plasmid. The yields were obtained from experiments performed in CGXII medium with 20 g/L of D-glucose and 20 g/L of D-fructose at 30°C and 130 rpm for 72 h (Fig. S5). (B) Production experiment of Fru^neg^-IolT1^G87S^Δ*zwf*::P_tuf_-*fbp*::GTG-*pgm* with the Mak expressing plasmid in CGXII medium with 20 g/L of D-glucose and 20 g/L of D-fructose for 72 h at 30°C and 130 rpm. In time intervals of 12 h, samples were taken. Growth was determined via OD_600_ measurement and sugar concentrations in the supernatant were determined via HPLC. The data points represent average values with standard deviations of three biological replicates.

For further improvement of the D-fructose 6-phosphate supply, its conversion to D-fructose 1,6-bisphosphate by phosphofructokinase PfkA should be reduced. As strong reduction of PfkA activity can lead to a complete loss of growth of *C. glutamicum* in D-glucose minimal medium (Liu et al., 2024), we avoided direct interventions with *pfkA* expression, e.g. via deletion, start codon exchange, or promotor exchange. Instead, we increased expression of the *fbp* gene for D-fructose 1,6-bisphosphatase (Fbp), which catalyzes the dephosphorylation of D-fructose 1,6-bisphosphate to D-fructose 6-phosphate, which is required for growth on substrates requiring gluconeogenesis, such as acetate, lactate or citrate (Rittmann et al., 2003). In a previous study, *fbp* expression was increased by a factor of 10 by exchanging its native promoter by the strong constitutive promoter of elongation factor TU (P_tuf_) (Becker et al., 2005).

By exchanging the native *fbp* promoter with the TU promoter in strain Fru^neg^-IolT1^G87S^Δ*zwf* slower growth, D-glucose and D-fructose consumption was observed (Fig. S5). The fructokinase-expressing strain showed a 21% increased final OD_600_ and a 45% decreased final D-fructose titer (4.36 ± 0.89 g/L) compared to the parental strain with the native *fbp* promoter (Fig. S5). For the xylose isomerase-expressing strain, such effects were not observed, as the final OD_600_ did not differ from the parental strain and final D-fructose titers were slightly increased (Fig. S5). Most importantly, expression of *fbp* by P_tuf_ had a strong positive effect on D-allulose production for both tested strains, reaching yields of 7.3 ± 0.4% and 7.0 ± 0.6% after 72 h for the fructokinase- and the xylose isomerase-based approach, respectively, which corresponds to increases of almost 50% and 37% compared to the precursor strain (Fig. 7A).

Besides the main branches of glycolysis and the pentose phosphate pathway, D-glucose 6-phosphate and D-fructose 6-phosphate can be further utilized for glycogen and GDP-mannose formation via the phosphoglucomutase (Pgm) and the mannose 6-phosphate isomerase (ManA), respectively (Fig. 6). Since deletion of *pgm* or *manA* can alter the cell morphology or impair the growth on D-glucose (Sasaki et al., 2011; Seibold and Eikmanns, 2013), expression of both genes was downregulated by a start codon exchange from ATG to GTG. While the start codon exchange of *manA* did not improve D-allulose production and resulted in similar yields for both fructokinase (6.8 ± 0.4%) and xylose isomerase-based approaches (6.7 ± 0.2%), the start codon exchange of *pgm* increased D-allulose formation for the fructokinase-based approach by 35% resulting in titers of 3.54 ± 0.04 g/L and a yield of 9.1 ± 0.1% (Fig. 7B). In contrast, the xylose isomerase-based approach did not reveal increased D-allulose formation upon GTG-*pgm* introduction and shows similar yields to the GTG-*manA* mutant (6.4 ± 0.3%) (Fig. 7A). The reason for this difference is still unclear. Interestingly, the fructokinase-carrying strain with GTG-*pgm* revealed a 28% higher D-fructose consumption compared to its parental strain and a 18% higher D-fructose consumption compared to its GTG-*manA* counterpart, which was not the case for the xylose isomerase-carrying strain whose D-fructose uptake remained almost unaltered compared to its parental strain (Fig. S5). In this regard, maybe the increased D-fructose uptake enables a more efficient supply of D-fructose 6-phosphate for Fru^neg^-IolT1^G87S^Δ*zwf*::P_tuf_-*fbp*::GTG-*pgm* and thus a higher formation of D-allulose.

## 4. Conclusion

This study examined the production of the low-calorie sweetener D-allulose by *C. glutamicum* in minimal medium using a pathway involving the epimerization of D-fructose 6-phosphate to D-allulose 6-phosphate followed by the dephosphorylation of D-allulose 6-phosphate, which should make this pathway irreversible. However, during construction and phenotyping of the production strains several challenges were encountered. In particular, the observed ability of fructokinase to phosphorylate D-allulose made the pathway, which was supposed to be irreversible, reversible again, causing degradation of previously formed D-allulose. As an alternative approach not requiring fructokinase activity at all, we used a recently evolved xylose isomerase that efficiently converts D-fructose and D-glucose at 30°C. This enables D-allulose production via D-glucose 6-phosphate formation and conversion to D-fructose 6-phosphate via D-glucose 6-phosphate isomerase. However, even in this case initially formed D-allulose was found to be degraded, presumably due to conversion to D-fructose by a side-activity of D-allulose 6-phosphate epimerase. By growth- or production-based enzyme and plasmid construct evaluation, we could not only identify the most suitable enzyme candidates for the process, but also determine the optimal order of the target genes on the pPREx2 expression plasmid to reach a comparably high conversion of D-glucose and D-fructose to D-allulose by the fructokinase- and the xylose isomerase-based pathways. Targeted metabolic interventions were applied to redirect carbon flux within the central metabolism of *C. glutamicum* towards enhanced D-allulose synthesis. Specifically, the deletion of *zwf* prevented D-glucose 6-phosphate oxidation in the pentose phosphate pathway, increased *fbp* expression improved the intracellular availability of D-fructose 6-phosphate, and modification of the *pgm* start codon should reduce D-glucose 6-phosphate–mediated glycogen formation. Together, these modifications effectively shifted metabolic flux toward D-allulose production, resulting in a 2.3-fold increase in yield. Eventually, a yield of 9.1% D-allulose was obtained from a D-glucose-D-fructose mixture with a fructokinase-based pathway. For increasing these titers, a variety of approaches can be followed, such as increasing the activity and specificity of D-allulose 6-phosphate epimerase and D-allulose 6-phosphate phosphatase and further reducing the flux of D-fructose 6-phosphate into glycolysis.

## Acknowledgements

This project was funded by the Bundesministerium für Bildung und Forschung (BMBF) within the project IMPRES-2 by a grant to M.B. (FKZ 031B1054B).

## Supplementary material

**Figure S1:**
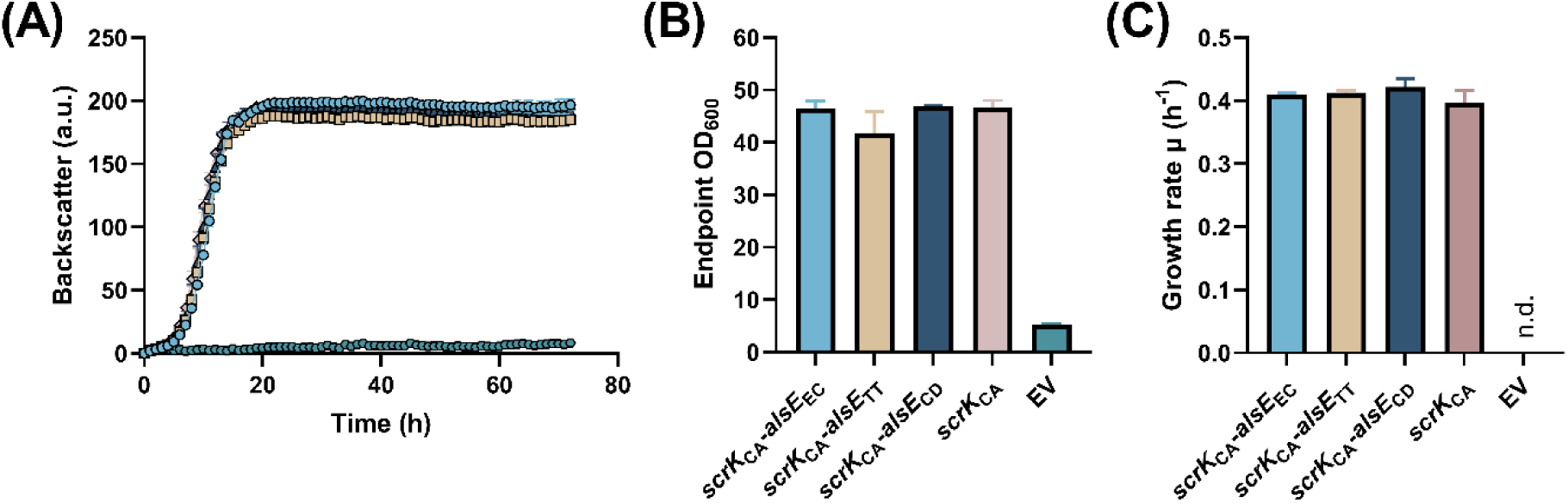
Impact of the fructokinase of *Clostridium acetobutylicum* (ScrK_CA_) alone and in combination with the allulose 6-phosphate 3-epimerase of *Escherichia coli* (AlsE_EC_), *Thermoanaerobacterium thermosaccharolyticum* (AlsE_TT_) or *Corynebacterium deserti* (AlsE_CD_) on the growth of *C. glutamicum* Fru^neg^-IolT1^T351P^ in fructose minimal medium. The corresponding genes were expressed from the *tac* promoter of the plasmid pPREx2. (A) Growth of the *C. glutamicum* strains in CGXII medium with 40 g/L of fructose at 30°C and 1200 rpm for 72 h. (B) Endpoint OD_600_ values of the strains after the growth experiment was terminated after 72 h. (C) Calculated growth rates of the respective strains in the period of 5-8 h. All data points given represent average values with standard deviations of three biological replicates. N.d., not determinable. EV, empty vector (pPREx2).

**Figure S2:**
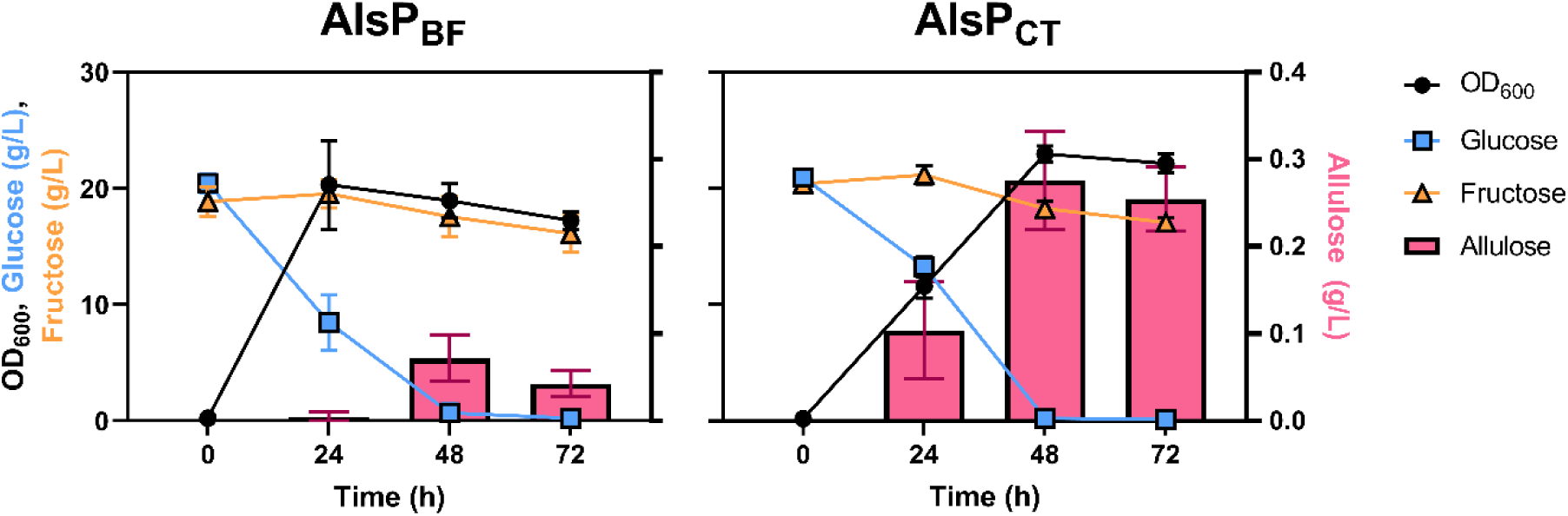
Screening of allulose 6-phosphate phosphatases from *Bacteroides fragilis* (AlsP_BF_) and *Clostridium thermocellum* (AlsP_CT_) for allulose formation in *C. glutamicum* Fru^neg^. The Fru^neg^ strain was transformed with pPREx2-*alsP*_BF_-*alsE*_EC_-*mak*_EC_ (AlsP_BF_) or pPREx2-*alsP*_CT_-*alsE*_EC_-*mak*_EC_ (AlsP_CT_) and cultivated in CGXII medium with 20 g/L of glucose and 20 g/L of fructose for 72 h at 30°C and 120 rpm. After 24, 48 and 72 h growth was measured via OD_600_ and the sugar content of the supernatant via HPLC. All data points given represent average values with standard deviations of three biological replicates.

**Figure S3:**
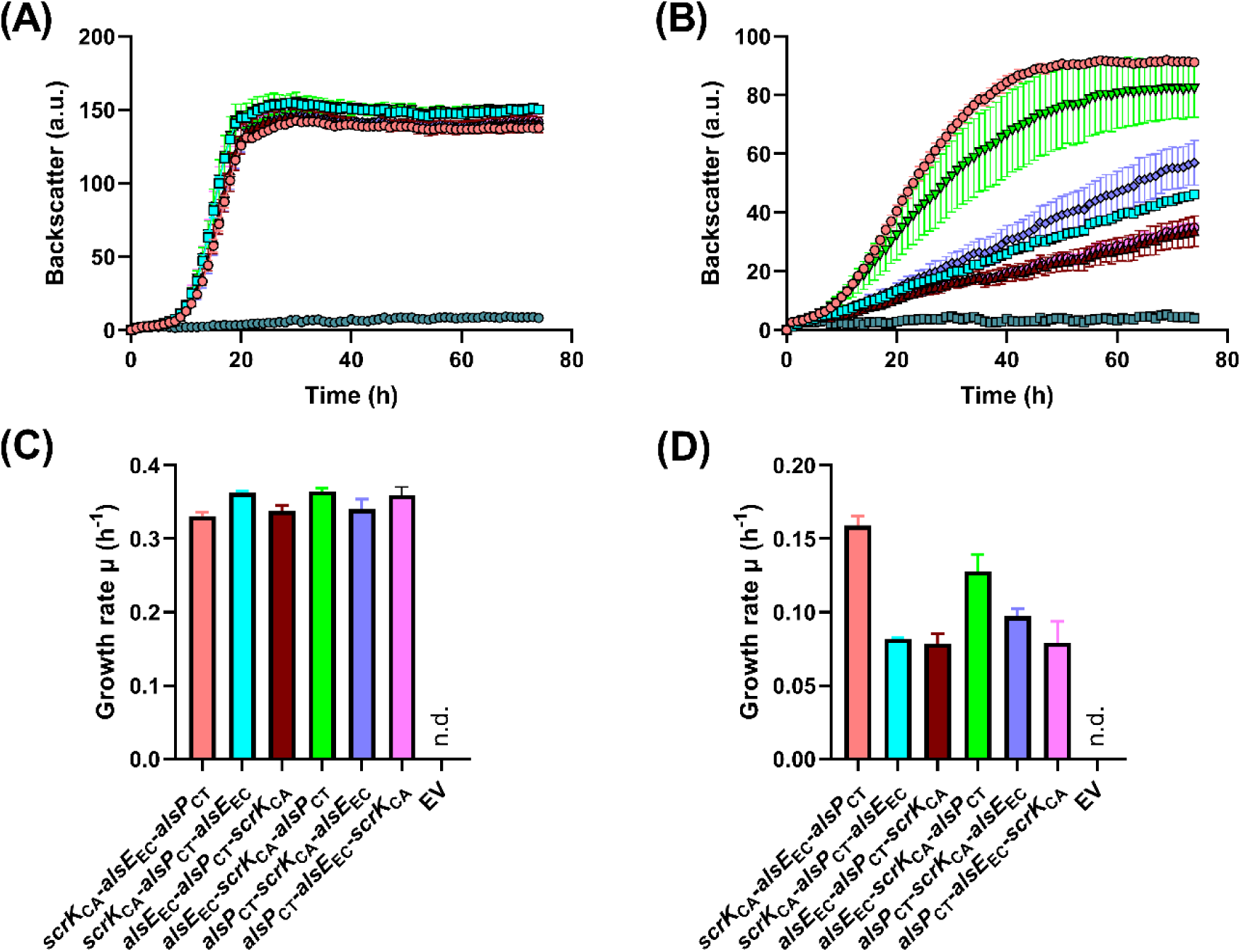
Plasmid-based gene shuffling experiment to identify the gene order which results in good growth on fructose minimal medium but poor growth on allulose minimal medium. (A, B) Growth experiment of *C. glutamicum* Fru^neg^-IolT1^T351P^ strains carrying the various pPREx2-based expression plasmids indicated in panels C and D in CGXII medium with 40 g/L of fructose (A) or 40 g/L of allulose (B). (C, D) Calculated growth rates of the respective strains on fructose (C) and allulose (D). The growth rate was calculated between 9 and 13 h for the strains cultivated on fructose and between 10 and 14 h for the strains cultivated on allulose. All data points represent mean values ± standard deviations from three biological replicates. N.d. not determinable.

**Figure S4:**
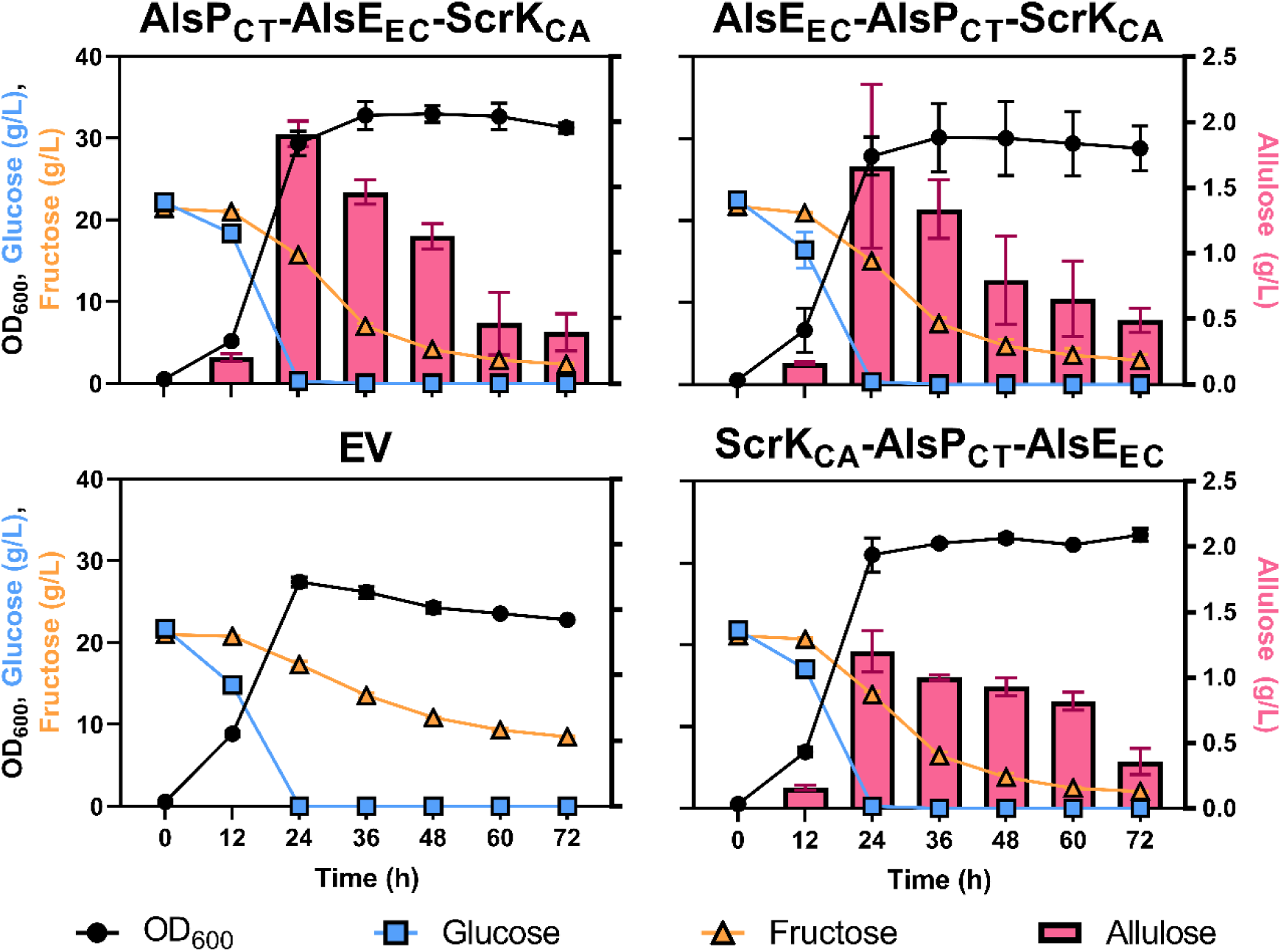
Investigation of allulose production by *C. glutamicum* Fru^neg^-IolT1^T351P^ with the growth-selected plasmids pPREx2-*alsP*_CT_-*alsE*_EC_-*scrK*_CA_ (AlsP_CT_-AlsE_EC_-ScrK_CA_), pPREx2-*alsE*_EC_-*alsP*_CT_-*scrK*_CA_ (AlsE_EC_-AlsP_CT_-ScrK_CA_), pPREx2-*scrK*_CA_-*alsP*_CT_-*alsE*_EC_ (ScrK_CA_-AlsP_CT_-AlsE_EC_) and the pPREx2 control (EV). The experiment was performed in CGXII medium with 20 g/L of glucose and 20 g/L of fructose at 30°C and 120 rpm for 72 h. At intervals of 12 h, samples were taken to measure growth via OD_600_ and sugar content of the supernatant via HPLC. All data points represent mean values ± standard deviations from three biological replicates.

**Figure S5:**
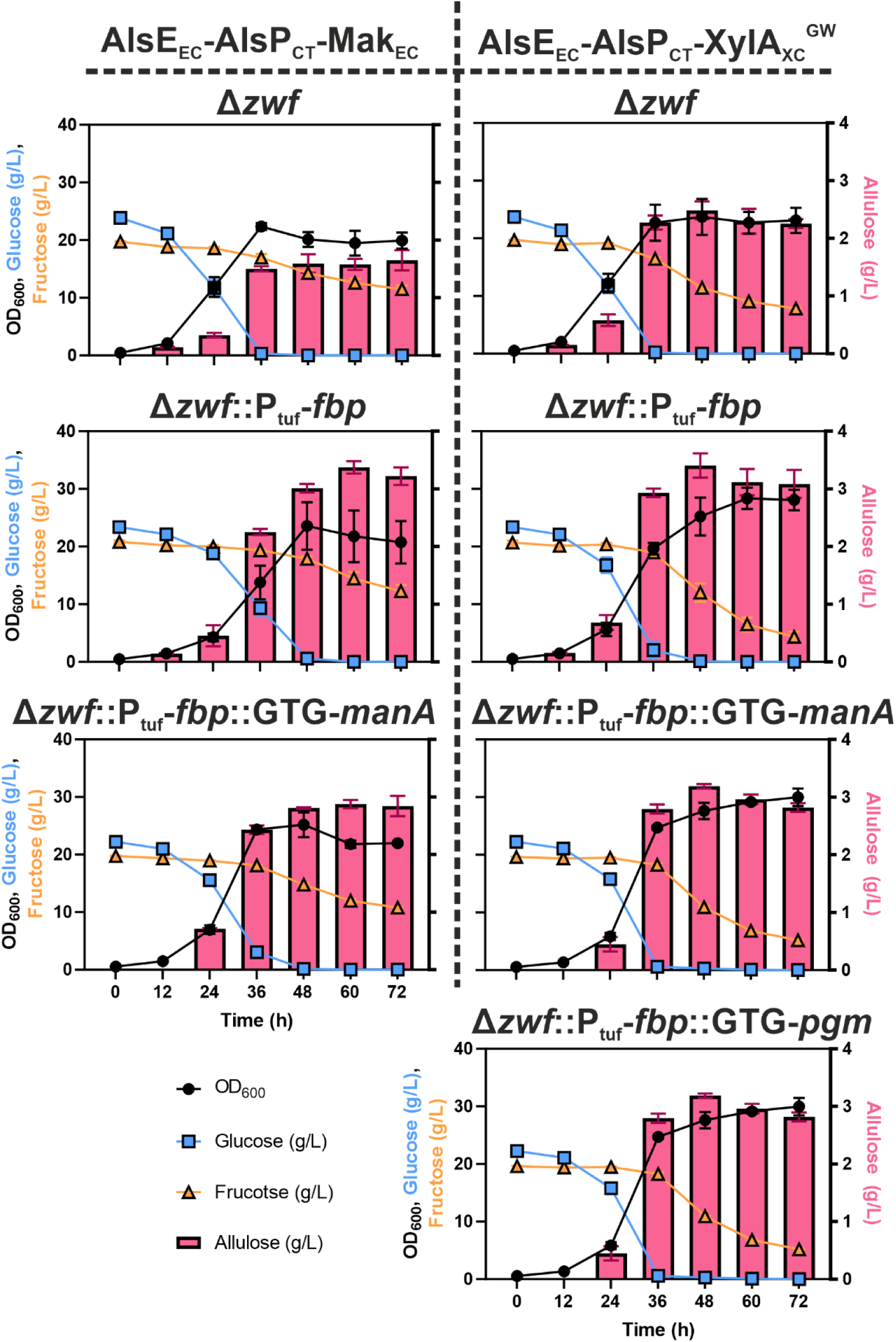
Analysis of allulose production for different *C. glutamicum* Fru^neg^-IolT1^G87S^ strains carrying pPREx2-*alsE*_EC_-*alsP*_CT_-*mak*_EC_ (AlsE_EC_-AlsP_CT_-Mak_EC_) or pPREx2-*alsE*_EC_-*alsP*_CT_-*xylA*_XC_^GW^ (AlsE_EC_-AlsP_CT_-XylA_XC_^GW^). For this experiment, different strains were constructed, which lack the glucose 6-phosphate dehydrogenase (Δ*zwf*), carry a *tuf* promotor exchange of the fructose 1,6-bisphosphatase (P_tuf_-*fbp*) and carry a start codon exchange of the mannose 6-phosphate isomerase (GTG-*manA*) or the phosphoglucomutase (GTG-*pgm*). The experiment was performed in CGXII medium with 20 g/L of glucose and 20 g/L of fructose at 30 °C and 120 rpm for 72 h. At intervals of 12 h, samples were taken to measure growth via OD_600_ and sugar content of the supernatant via HPLC. All data points represent mean values ± standard deviations from three biological replicates.

